# Plant-derived, nodule-specific cysteine rich peptides inhibit growth and psyllid acquisition of ‘*Candidatus* Liberibacter asiaticus’, the citrus Huanglongbing bacterium

**DOI:** 10.1101/2023.06.18.545457

**Authors:** Steven A. Higgins, David O. Igwe, John S. Ramsey, Stacy L. DeBlasio, Marco Pitino, Randall Niedz, Robert G. Shatters, Laura A. Fleites, Michelle Heck

## Abstract

The Asian citrus psyllid, *Diaphorina citri*, is a vector of ‘*Candidatus* Liberibacter asiaticus’ (*C*Las), a gram-negative, obligate biotroph whose infection in *Citrus* species is associated with citrus greening disease, or Huanglongbing (HLB). Strategies to block *C*Las transmission by *D. citri* remain the best way to prevent the spread of the disease into new citrus growing regions. However, identifying control strategies to block HLB transmission poses significant challenges, such as the discovery and delivery of antimicrobial compounds targeting the bacterium and overcoming consumer hesitancy towards accepting the treatment. Here, we computationally identified and tested a series of 20-mer nodule-specific cysteine-rich peptides (NCRs) derived from the Mediterranean legume, *Medicago truncatula* Gaertn. (barrelclover) to identify those peptides that could effectively prevent or reduce *C*Las infection in citrus leaves and/or prevent *C*Las acquisition by the bacterium’s insect vector, *D. citri*. A set of NCR peptides were tested in a screening pipeline involving three distinct assays: a bacterial culture assay, a *C*Las-infected excised citrus leaf assay, and a *C*Las-infected nymph acquisition assay that included *D. citri* nymphs, the only stage of *D. citri*’s life-cycle that can acquire *C*Las leading to the development of vector competent adult insects. We demonstrate that a subset of *M. truncatula*-derived NCRs inhibit both *C*Las growth in citrus leaves and *C*Las acquisition by *D. citri* from *C*Las-infected leaves. These findings reveal NCR peptides as a new class and source of biopesticide molecules to control *C*Las for the prevention and/or treatment of HLB.

## Introduction

Huanglongbing (HLB), also known as citrus greening disease, is currently the most devastating disease affecting citrus production worldwide (McCollum and Baldwin 2016). The causal bacteria of HLB are phloem-limited, gram-negative *Alphaproteobacteria* in the genus *Liberibacter* and includes: ‘*Candidatus* Liberibacter asiaticus’ (CLas), found in Asia, North and South America, Oceania and the Arabian Peninsula (Bove 2006; Haapalainen 2014), ‘*Ca*.

Liberibacter americanus’ (*C*Lam), found in South America (Teixeira et al. 2008; Texeira et al. 2005), and ‘*Ca*. Liberibacter africanus (*C*Laf) found in Africa and the Arabian Peninsula (Garnier and Bové 1996; Pietersen et al. 2010). The natural mode of spread for these three bacteria is by insect vectors (Ammar et al. 2016; da Graca et al. 2016; Hall et al. 2013; Huang et al. 2020). Two insect vectors of HLB-associated *Ca*. Liberibacter spp. have been identified: the Asian citrus psyllid, *Diaphorina citri* Kuwayama (Hemiptera: Liviidae) (Bove 2006; Capoor et al. 1967) (Capoor *et al*., 1967; Bové, 2006), and the African citrus psyllid, *Trioza erytreae* del Guercio (Hemiptera: Triozidae) (Van den Berg et al. 1992). According to the database maintained by the European and Mediterranean Plant Protection Organization (EPPO, https://gd.eppo.int/), *D. citri* possesses the widest geographical range of the two *C*Las insect vectors, which may partially explain why *C*Las is the most wide-spread bacterium related to HLB worldwide. Other explanations may also include difference in virulence or insect transmissibility among the different species of Liberibacter.

*C*Las infection is confined to the citrus vascular tissue, specifically the phloem (Huang et al. 2020). This tissue tropism in the plant facilitates transmission by *D. citri*, a hemipteran insect with piercing sucking mouthparts to feed on the phloem sap, and also demands that HLB antimicrobial therapies be adequately delivered to and disseminated systemically within phloem tissue. *C*Las transmission by *D. citri* is circulative and propagative, meaning the bacterium must circulate throughout the insect vector’s body and infect insect tissues through active replication (Ammar et al. 2016; Higgins et al. 2022; Inoue et al. 2009). Following insect feeding on an infected tree, *C*Las must cross the insect’s epithelial gut barrier, traverse through the hemolymph, and enter into the salivary glands prior to transmission to a new tree (Ammar et al. 2011a; Ammar et al. 2016; Ammar et al. 2011b; Inoue et al. 2009). Transmission of *C*Las by *D. citri* adults is more efficient when bacteria are acquired in the insect’s juvenile stage (Ammar el et al. 2016; Inoue et al. 2009), where *C*Las acquisition is facilitated by a combination of long periods of phloem ingestion and significant immune suppression that occurs during the nymphal stage (Mann et al. 2018; Ramsey et al. 2017). Lee *et al*., (2015) proposed the idea of ‘flush transmission’ of *C*Las from infected *D. citri* adults to their young nymphs, when these nymphs feed on the flush leaves recently infected by their parents (Lee et al. 2015). ‘Flush’ is any new leaf growth ranging in development from first emergence up until the leaves are fully expanded yet still tender (Hall et al. 2016). Young flush becomes infectious within 10-15 d after receiving *C*Las inoculum from infected *D. citri* (Lee et al. 2015). Developing nymphs acquire *C*Las from these infected feeding sites and emerge as vector competent adults immediately capable of spreading *C*Las to a new flush point, repeating the process. The bacteria replicate to high levels in the psyllid salivary glands by two weeks after adult emergence (Ammar et al. 2016). Thus, new antimicrobial therapies that are phloem-mobile and can block *C*Las acquisition by nymphs are paramount for the control of HLB (Kennedy et al. 2023).

*C*Las is genetically related to *Sinorhizobium meliloti*, an economically important nitrogen-fixing plant symbiotic bacterium (Kuykendall et al. 2012). *S. meliloti* infects the roots of leguminous plants like alfalfa, triggering the formation of a specialized plant organ (nodule) that is colonized by the bacterium. Plant expression of a large diversity nodule-specific cysteine-rich peptides (NCRs) that govern *S. meliloti*’s replication and functioning to transform the cells into a nitrogen-fixing, symbiotic form (Tiricz et al. 2013b). NCRs are similar in sequence to defensins, a group of bactericidal antimicrobial peptides (AMPs) that function to kill microbes as part of the immune response in plants and insects (Broekaert et al. 1995; Mergaert et al. 2003). Notably, two of these NCRs have been demonstrated to act as broad-spectrum bactericides of plant and human bacterial pathogens by disrupting their cell membrane (Mikulass et al. 2016), but the antimicrobial potential of other NCRs remains unexplored. The combination of large functional diversity, high stability in nature (Durgo et al. 2015), ability to kill genetically diverse bacterial species (Mikulass et al. 2016), and a simple one-gene, one-product expression system, make NCRs an excellent, untapped strategy for the control of HLB. In this work, we computationally predicted NCR peptides from the *M. truncatula* genome. We developed a multi-tiered screening approach using both plants and insects to identify NCR peptides that demonstrate antimicrobial activity against *C*Las. A subset of five NCR peptides inhibited *C*Las growth in leaves and prevented the development of high titer, vector-competent *D. citri* adults most likely to spread *C*Las within a grove.

## Materials and methods

### Plant and insect growth conditions

*Diaphorina citri* Kuwayama (Hemiptera: Liviidae) were reared in controlled growth chambers in Ithaca, NY on *Citrus medica* (citron) plants under 14:10 h light:dark cycle at 28 °C. The colony-supporting citrus plants were regularly monitored by lab technicians, maintained by pruning and watered on a weekly or as needed basis and fertilized bi- weekly. Individual *D. citri* nymphs from citron colonies were manually removed using a fine paintbrush prior to transfer to *C*Las infected excised leaves for the excised leaf acquisition assay as described in (Igwe et al. 2022). *C*Las-infected citron plants were reared separately from healthy citron under the same photoperiod and growth conditions. Infected plants were generated using *D. citri* inoculation and monitored for HLB development by periodic analysis of *C*Las 16S rDNA gene copies using quantitative PCR (qPCR) and observation of symptom development.

### Computational prediction of 20-mer NCR peptides from the *M. truncatula* genome for peptide synthesis

A database of 662 *M. truncatula* nodule-specific cysteine rich (NCR) peptide sequences was compiled by searching the annotated *M. truncatula* proteome in UniProt Knowledgebase (UniProtKB) using a keyword search (*e.g.* NCR, nodule cysteine-rich). The Random Forest algorithm antimicrobial peptide (AMP) prediction tool on CAMPR3 (http://www.camp.bicnirrh.res.in/predict_c/) was used to identify 20-mer peptide sequences with antimicrobial characteristics within each of these 662 proteins (doi:10.1093/nar/gkp1021). Out of the 662 proteins identified, at least one 20mer peptide sequence with an AMP score >0.5 was identified for 623 proteins. The grand average of hydropathicity index (GRAVY) score for each peptide was predicted using the method of Kyte and Doolittle (Kyte and Doolittle 1982).

GRAVY scores calculate the sum of the hydropathy values of all the amino acids in the peptide divided by the sequence length. The GRAVY score was calculated for peptides with an AMP score greater than 0.5 (https://www.gravy-calculator.de/index.php). Negative peptide GRAVY scores typically predict peptides will be water-soluble, a desirable characteristic for facilitating delivery to plants and for testing in cells. However, some known antimicrobial peptides have positive GRAVY scores, so a range of GRAVY scores were considered. A total of 182 peptides with a range of GRAVY scores (163 negative, 20 positive) were selected for synthesis and prioritized for screening based on these and other characteristics, such as whether they had been previously localized to the bacteroid or shown to have AMP activity in other papers (Table 1, Table S1). Any secretion signals, which are not required for antimicrobial activity, were removed prior to synthesis (Tiricz et al. 2013a). Sequences were synthesized by Biomatik USA, LLC (Wilmington, DE, USA) to between 70 and 98% purity according to their standard synthesis protocols, subjected to removal of the trifluoroacetic acid and sequence validated using mass spectrometry and high-pressure liquid chromatography by the company prior to shipment to the lab.

**Table 1.** Annotated NCR sequences in UniProt

|  | Length | Number of NCRs | Percentage of NCRs |
| --- | --- | --- | --- |
| Tier 1<sup>a</sup> | ≤30 | 14 | 2.32% |
| Tier 2<sup>b</sup> | 31-60 | 232 | 38.41% |
| Tier 3<sup>c</sup> | 61-90 | 323 | 53.48% |
| Tier 4<sup>d</sup> | >91 | 35 | 5.79% |

<sup>a</sup>Tier 1: peptides with functional validation and/or verified localization to nodule or bacteroid tissue, and peptides without validation that are 30 residues or smaller.
<sup>b</sup>Tier 2: peptides 31-60 residues in length, with higher priority given to sequences with antimicrobial characteristics (multiple cysteine residues and/or overall cationic character).
<sup>c</sup>Tier 3: peptides 61-90 residues in length, with higher priority given to sequences with antimicrobial characteristics (multiple cysteine residues and/or overall cationic character).
<sup>d</sup>Tier 4: peptides that are greater than 91 residues in length, with lowest priority for screening.

### Bacterial culture assays

Growth of *Liberibacter crescens* strain BT-1 was performed in BM7 basal salts (BM7) medium. The BM7 basal salts solution was prepared by combining and dissolving 2 g alpha ketoglutaric acid sodium salt, 10 g ACES buffer, and 3.75 g potassium hydroxide in distilled water, tuning the pH to 6.9, and adjusting the final volume to 550 ml. After autoclaving and cooling to RT, fetal bovine serum (FBS, 150 ml) and TMN-FH medium (300 ml) were added aseptically to the BM7 basal salts solution by stirring. Agar medium was prepared similarly, except that 15 g of microbiological grade agar was added to the basal salts solution prior to autoclaving, and FBS and TMN-FH medium were pre-warmed in a water bath and added at 50 °C instead of RT. Prior to preparation of growth assays, *L. crescens* strain BT-1 was streak plated from glycerol freezer stocks onto sterile BM7 basal salts (hereafter BM7) agar medium and allowed to grow at 28 °C for 7 to 14 days, at which point a single colony was picked and inoculated into 3.5 ml BM7 broth and grown at 28 °C with shaking at 200 rpm. An aliquot of this culture (5 % v/v inoculum) was transferred to fresh BM7 broth and allowed to grow to an O.D. 600nm of 0.4 to 0.7 (roughly 3 to 4 days). The transfer culture was then diluted to an O.D. 600nm of 0.025, and 50 microliters of diluted culture were combined in a sterile low-bind, round bottom polypropylene plate (Costar 3879) with 50 microliters of NCR peptide (2 or 0.2 mg/ml) diluted in BM7 medium. The final concentrations of NCR peptides in all assays were 1 or 0.1 mg/ml. The antimicrobial peptide polymyxin B sulfate (hereafter PMB; 0.5 mg/ml final concentration) dissolved in BM7 broth, *L. crescens* strain BT-1 cells in BM7 broth, and BM7 broth without cells were utilized in every 96-well plate assay as positive growth inhibition, no growth inhibition, and no growth controls, respectively. The growth inhibition assays for all NCR peptides examined were performed in at least duplicate over multiple preparations of *L. crescens* cells. Following their preparation, 96-well plates were wrapped with parafilm, loaded into a small plastic container lined with moist paper towels, and incubated at 28 °C with shaking (200 rpm) over a seven-day period. The O.D. 600 nm was recorded daily on a Synergy HT plate reader (Agilent Technologies, Inc., Santa Clara, CA, USA).

### Excised leaf assay

*C*Las-infected plants were first screened for infection using a standard *C*Las 16S rDNA qPCR assay with nucleic acids extracted from several small leaf discs dissected from at least N = 6 leaves per plant. Mature leaves with the petiole intact were then selected for the excised leaf assays from plants identified as having reliably detectable *C*Las 16S rDNA gene copies in the excised leaves selected (Hosseinzadeh et al. 2019; Li et al. 2006). The petiole was then submerged in solutions of 0.1 mM potassium phosphate buffer, pH 5.8, (hereafter KPO4 buffer) containing either 1 mg/ml PMB or 1 mg/ml of each NCR peptide. All leaves (n = 10 per treatment combined from multiple infected trees) and solutions were housed in a 0.2 ml sterile PCR tube and wrapped with parafilm to prevent evaporation of the solution. After an overnight incubation period in which most of the solution was taken up by the excised leaf, the PCR tube with leaf was inverted, flicked gently to clear any liquid from the tip of the PCR tube so as to ensure any remaining NCR peptide in solution would not be discarded, and the bottom of the PCR tube was cut with 70% (v/v) ethanol-wiped scissors. The leaf petiole and PCR tube were then submerged inside a sterile 2 ml microcentrifuge tube containing KPO4 buffer where they remained for 7 days to complete uptake of any remaining NCR peptide and incubate the *C*Las- infected excised leaf (**Fig. S1**). The KPO4 buffer was topped off as needed, usually on a daily basis and leaves were maintained in a 70 °F chamber with 14 h light:10 h dark schedule. A set of four leaf discs (1 mm diameter) were extracted from an excised leaf at 0 and 7 days post access to all treatment solutions, added to a 2 ml microcentrifuge tube, and snap frozen in liquid nitrogen. All samples were stored at -80 ℃ until nucleic acid extractions were performed. **Nymph acquisition assay**. To test whether the NCR peptides had an impact on *C*Las acquisition by *D. citri,* we performed a psyllid nymph acquisition assay using *D. citri* (Igwe et al. 2022). As above, we evaluated *C*Las titers in citrus colonies using a single leaf punch from multiple colony leaves to determine which plants are useful to harvest leaves for the nymph acquisition assays.

The *C*Las-infected plants used to provide leaves for this assay were psyllid-free at the time of sample collection for DNA extraction and qPCR analysis. For the NCR peptide assays, plants with Cq < 30 were used to supply the excised leaves for the analysis. Leaves from *C*Las-infected with Cq values of 26.11, 27.37 and 28.02 were selected for these experiments. Branch stems from these *C*Las-infected citron plants were collected and cut at a 45° angle with some portion of the stem still intact to facilitate water absorption via the exposed cut surface area of the citron stem supporting a single leaf. Cuttings were transferred to sterile 0.6 mL tubes containing 200 µL NCR peptides (803543 and 803570, 20 mg/ml) diluted with KPO4 buffer to 1 mg/mL, and PMB (0.5 mg/mL) in KPO4 buffer. All cuttings were incubated for 24 hours to allow adequate uptake of the solutions then transferred to 2.0 mL tubes, wrapped with parafilm, and placed inside 50 mL falcon tubes. The prepared *C*Las-infected citrus leaves from three infected plants were distributed equally among the treatments. The lids of the falcon tubes were modified to contain a mesh screen to enhance ventilation. Healthy psyllid nymphs (*C*Las-free) of 2^nd^ and 3^rd^ instars were collected from *D. citri* colonies reared on healthy citron plants and transferred to a plastic weigh boat to remove accumulated honeydew using a fine-tipped paint brush. Nymphs were starved for two hours before transferring them to a single citrus leaf incubated in one of the treatment solutions described above. To each leaf we added a total of 10 psyllid nymphs and each treatment was comprised of 10 individual excised citrus leaves (n = 100 psyllid nymphs per treatment). The assay buffers were monitored and their volume maintained by replacement with water for 21 days, after which the adult psyllids were collected for DNA extraction and qPCR analysis.

### Plant and insect DNA extractions

Total nucleic acids were extracted from individual adult psyllids and citrus leaves (three punches per leaf) as described in (Igwe et al. 2022). All nucleic acid extracts were quantified using a NanoDrop 2000 spectrophotometer (Thermo Fisher Scientific, Inc., Waltham, MA, USA) and stored at -80 °C until analysis.

### Detection and quantification of *C*Las rDNA and rRNA in plants and insects

For the detection of *C*Las 16S rDNA genes, qPCR assays were performed on an Applied Biosystems QuantStudio 6 Flex Real-Time PCR System (Thermo Fisher Scientific, Waltham, MA, USA). The TaqMan Universal PCR master mix (Thermo Fisher Scientific, Waltham, MA, USA) with 16S rRNA gene primer and probe sets (Hosseinzadeh et al. 2019; Li et al. 2006; Ramsey et al. 2015) were used. Briefly, the *C*Las 16S rDNA gene primers used were *C*Las16SF (5’- TCGAGCGCGTATGCAATACG-3’), *C*Las16SR (5’-GCGTTATCCCGTAGAAAAAGGTAG-3’), and probe *C*Las16Sp (5’-AGACGGGTGAGTAACGCG-3’). *C*Las titers in all samples were quantified in duplicate for each biological replicate. The final concentration of qPCR mix contained 1X TaqMan Universal PCR master mix (10 µL) (Thermo Fisher Scientific, Inc.), 1 µL of each of forward and reverse primers (10 µM each), 1 µL of probe (5 µM), 2 µL of 25 ng/µL DNA sample, and 5 µL of nuclease free water in a total volume of 20 µL. The qPCR program consisted of 2 min incubation at 50 °C followed by 10 min incubation at 95 °C and 40 cycles at 95 °C for 15 s and 60 °C for 1 min. For the analysis of *C*Las 16S rRNA genes, nucleic acid extracts were split into two equal portions, one of which was subjected to DNase I treatment using the TURBO DNA-free kit (Thermo Fisher Scientific, Inc.). Briefly, each reaction contained 2.5 µL of 10X DNase I buffer, 0.5 µL of DNase I enzyme, and 17 µL of nucleic acid extract. DNase reactions were incubated at 37 °C for 1 hr, followed by addition of 5 µL of DNase inactivation reagent and incubation at RT for 5 min, followed by centrifugation for 5 min at 2,000 x g. Following DNase I treatment, 15 µL of the supernatant above the dense inactivation reagent was transferred to a fresh, sterile DNase/RNase-free 96-well plate for first-strand cDNA synthesis. The iScript cDNA synthesis kit (Bio-Rad Laboratories, Inc., Hercules, CA, USA) was used to generate the first-strand cDNA by combining 15 µL DNase I treated RNA with 4 µL of 5X iScript master mix reagent, 0.5 µL of iScript reverse transcriptase, and 0.5 µL of DNase/RNase-free water. Samples were vortexed and centrifuged briefly and then incubated on a MiniAmp thermal cycler (Applied Biosystems) using a thermal program consisting of 25 °C for 5 min, 46 °C for 40 min, 95 °C for 1 min, and 4 °C thereafter. All first-strand cDNA and RT-minus RNA controls were stored at –80 °C until qPCR analysis. We calculate and report “CLas cell equivalents” (16S rDNA gene copies/3) as reported in Igwe et al. (2022). All qPCR standard curve information used to calculate gene copy numbers can be found in Supplementary Information **Table S2**.

### Statistical analysis

All experiments were performed with replication and included a minimum of technical duplicates (cell plate growth assays) and between six and ten replicate leaves for detached leaf and nymph acquisition assays, respectively). Statistical analyses visualizations were performed in R v4.1.2 and RStudio v2021.09.1+372 (RStudio 2020) using packages tidyverse v1.3.1 (Wickham et al. 2019), vegan v2.6.2 (Oksanen et al. 2013), growthrates v0.8.3 (Petzoldt 2022), ggridges v0.5.3 (Wilke 2021), grid v4.1.2 and gridExtra v2.3 (Auguie and Antonov 2017; RStudio 2020), cowplot v1.1.1 (Wilke 2020), extrafont v0.18 (Winston 2022), stringr v1.4.0 (RStudio 2020; Wickham 2019), readxl v1.3.1 (Wickham and Bryan 2019), ggpubr v0.4.0 (Kassambara 2020), ggbiplot v0.55 (Vu 2011), ggrepel v0.9.1 (Kamil 2021), and FrF2 v2.2.3 (Grömping 2014). The growth rates of *L. crescens* strain BT-1 were estimated using the growthrates R package by fitting a logistic growth model to optical density measurements collected over time from *L. crescens* strain BT-1 cells grown in a 96-well plate. Starting parameters provided to the growth model were estimated from the total data set and included parameters y0, mumax, and K, which are the intercept, maximum growth rate, and carrying capacity of the population, respectively. Lower and upper bound estimates for each parameter were also estimated from the complete data set and provided to the model (see R code in SI for details). Our resolution five fractional factorial assay design was generated using the VF2 algorithm implemented in the FrF2 R package (Cordella et al. 2001). The fractional factorial design permitted exploration of two factor interactions for ten NCR peptides (at both 0.1 and 1 mg/ml concentrations) by assaying the effects of combinations of NCR peptides on the growth of *L. crescens* strain BT-1 without performing all pairwise comparisons. Thus, a total of 1,023 pairwise NCR peptide combinations from ten peptides were reduced to 128 NCR peptide combinations for testing in 96-well plate growth assays by the fractional factorial design. Results from the fractional factorial design assays were tested for significance (α < 0.05) using a two- way ANOVA analysis in R and main and interaction effects plots were generated using the MEPlot and IAPlot functions in the FrF2 package. We performed paired Welch’s t-tests with unequal variance among groups and Wilcoxon rank sum tests in R to identify statistically significant (α < 0.05) changes in log-transformed mean 16S rRNA gene copies or the ratio of 16S rRNA to rDNA gene copies between the two time points examined. A principal components analysis was performed using the prcomp function in R to examine the relationship between physicochemical properties of the NCR peptides and their growth inhibition of *L. crescens* strain BT-1 in 96-well plate assays. All data were centered and scaled prior to performing the analysis. Spearman rank correlations were performed using the cor.test function in R. Welch’s analysis of variance followed by a Dunnet’s test was used to compare NCR peptide treatments to the buffer control in the nymph acquisition assays. For the analysis of psyllid mortality following NCR peptide treatments, Shapiro-Wilk normality test (Shapiro and Wilk 1965), and Levene’s test of homogeneity of variance (Levene 1960) were performed on the number of dead psyllid nymphs per treatment with the NCR peptides, PMB and KPO4 buffer control. The data were not normally distributed, and there was equal variance. Based on the nature of the distribution and existing equal variance in the data, Kruskal-Wallis variance analysis (a non-parametric analysis of variance) method was used to test for treatment effects (Kruskal and Wallis 1952).

## Results

### Computational analysis of NCR peptides identifies a subset of 20-mers for synthesis

Our analysis revealed a total of 662 *M. truncatula* proteins annotated as NCRs in UniProt. These peptides ranged from 16 to 924 residues in length, with the majority being 90 residues or smaller (**Table 1**). Peptides were categorized for screening based on a set of parameters we developed. A total of 14 Tier 1 peptides included both peptides with previous functional validation and/or verified localization to nodule or bacteroid tissue in the literature and peptides without validation that were 30 residues or smaller. More than 230 Tier 2 peptides were identified and included those 31-60 amino acid residues in length, with higher priority given to sequences with antimicrobial characteristics (multiple cysteine residues and/or overall cationic character). The majority of the peptides, 323 in total, were Tier 3 peptides and included those 61-90 residues in length with higher priority for screening given to sequences with antimicrobial characteristics. A total of 183 smaller, 20-mer antimicrobial peptides were computationally derived from the larger sequences for synthesis based on CAPM prediction and their GRAVY scores (**Table S1**). These peptides were selected based on their GRAVY scores.

### Bacterial culture assay reveals NCR peptide 20-mers have a range of activities against *L. crescens*, a cultivable surrogate of *C*Las

We performed a high-throughput *in vitro* antimicrobial assay to test the effect of 183 NCR peptides for growth inhibition of *Liberibacter crescens* strain BT-1, a cultivable surrogate of ‘*Candidatus* Liberibacter asiaticus’ (Jain et al. 2019). While the individual NCR peptides had a wide range of effects on the growth of *L. crescens* strain BT-1 including growth promotion (**Fig. 1, purple distribution**), among the peptides that had an effect, more peptides showed growth inhibitory effects (**Fig. 1, teal distribution**). Note, the negative control in this case is growth of *L. crescens* strain BT-1 without any inhibitory molecule (**Fig. 1, green distribution**). Overall, the NCR peptides were observed to inhibit the growth rate of *L. crescens* strain BT-1 by up to 73% (**Fig. 2**), but some peptides resulted in an increase in *L. crescens* growth. An analysis of the physiochemical properties of the NCR peptides revealed weak but positive correlations between the *L. crescens* growth rate inhibition and number of cysteine residues, net peptide charge at pH 7 and predicted peptide solubility as a function of the GRAVY score (**Fig. S2**).

**Figure 1.**
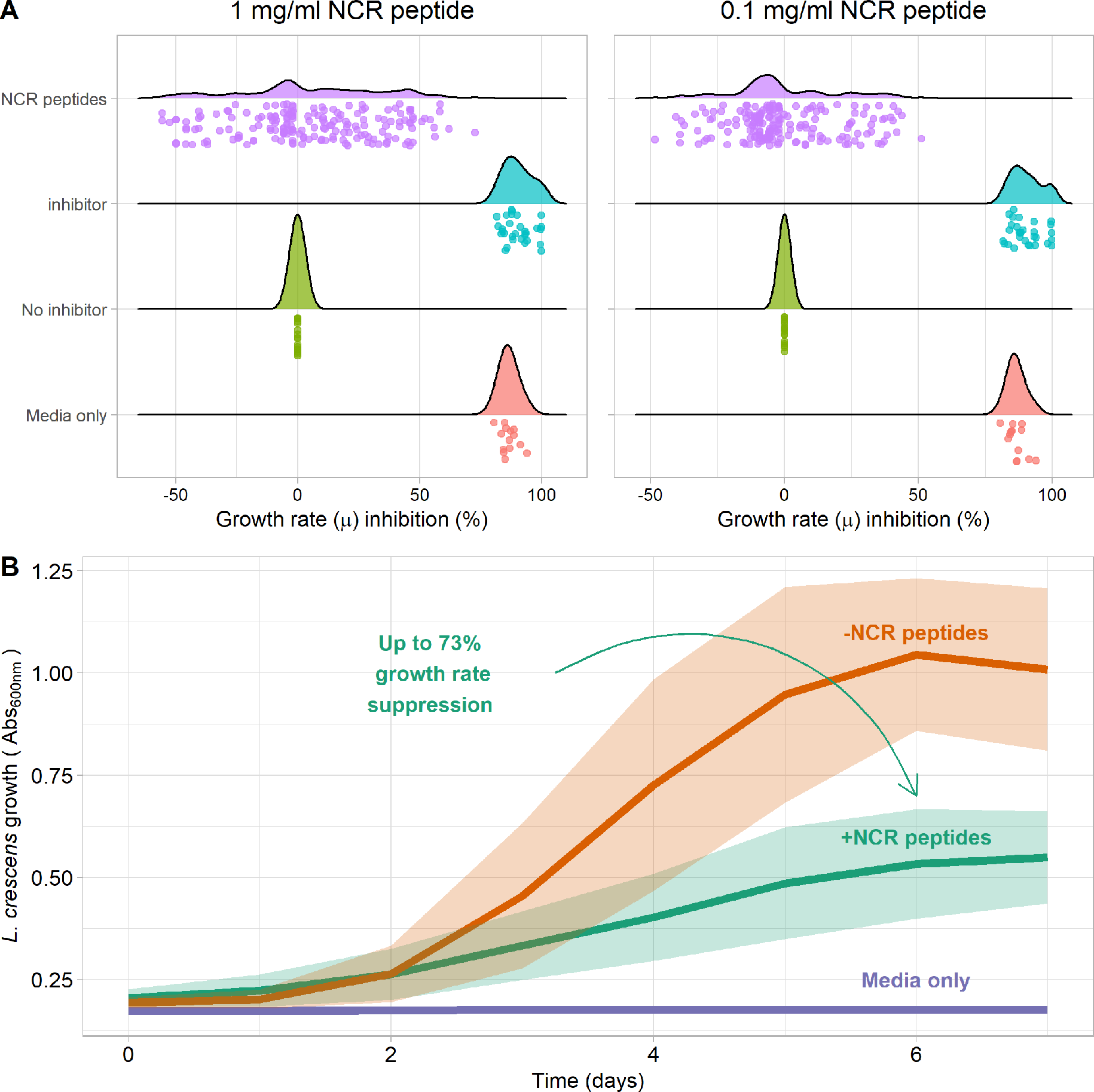
Growth rate inhibition of *Liberibacter crescens* strain BT-1 in BM7 medium by NCR peptides. We present the distribution of growth rate inhibition observed for individual NCR peptides at 0.1 and 1.0 mg/ml concentrations (A) and the altered growth curve for combined 1 mg/ml NCR peptides compared to no peptide controls and uninoculated medium (B).‘NCR peptides’ refers to the 182 NCR peptides supplied in the medium at a final concentration of 1 or 0.1 mg/ml. The row ‘inhibitor’ refers to the antimicrobial peptide polymyxin B sulfate (0.5 mg/ml). The ‘No inhibitor’ row refers to the growth of *L. crescen*s strain BT-1 in BM7 medium without inhibitors. The ‘Media only’ is BM7 medium only blank control. All values on the x-axis are percent growth rate inhibition, calculated as (‘No inhibitor’ growth rate (µ) - growth rate (µ) / ‘No inhibitor’ growth rate (µ)) * 100). Hence, when the x-axis value is negative, the maximum estimated growth rate of *L. crescens* BT-1 was greater than the maximum estimated growth rate of the no inhibitor added control. Note, the logistic growth models applied to the data usually provided non-zero maximum growth rate estimates for ‘inhibitor’ and ‘Media only’ treatments although their change in absorbance over time was zero. Hence, the growth rate inhibition on the x-axis is not always 100%.

**Figure 2.**
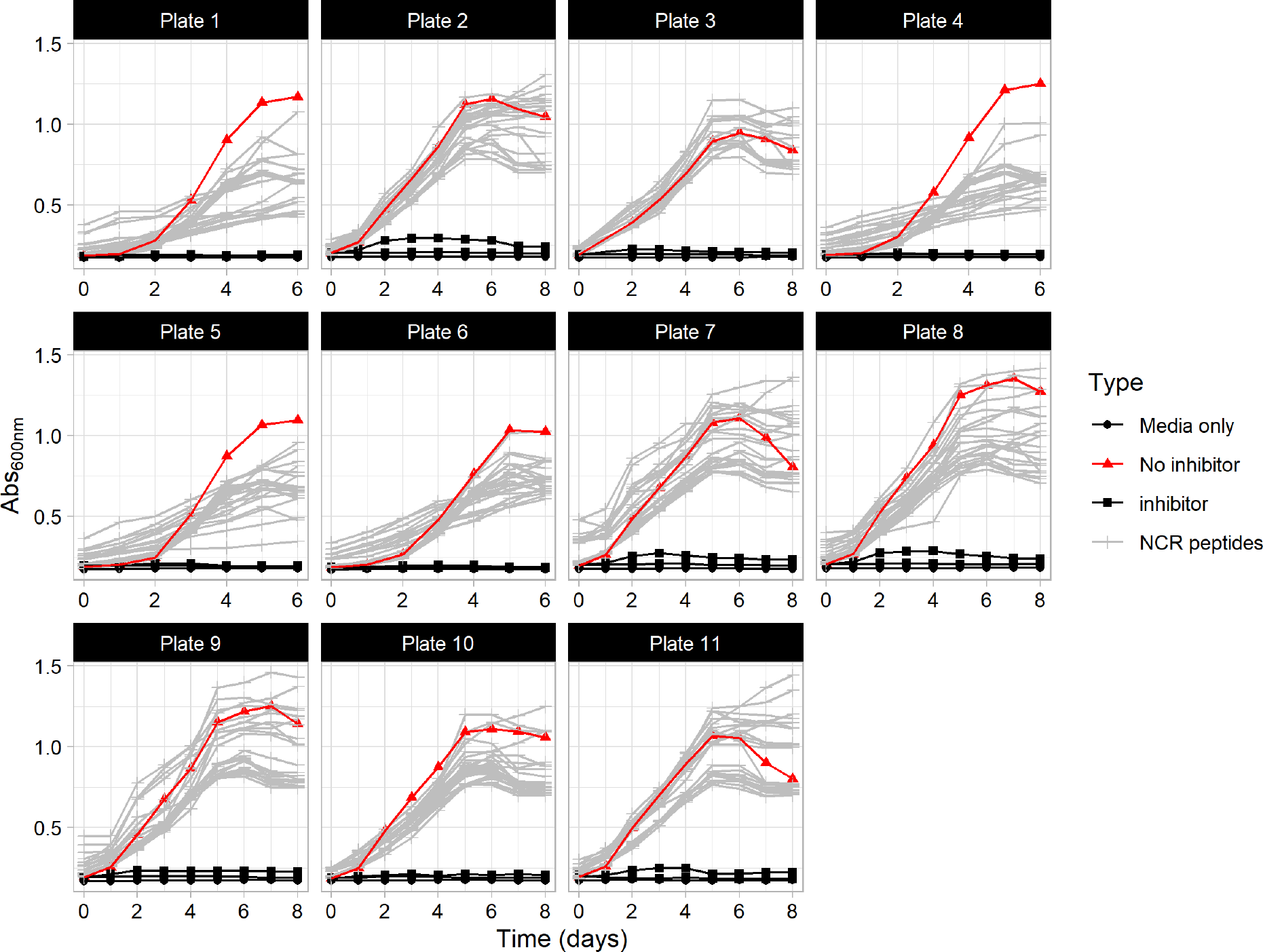
Growth inhibition of *Liberibacter crescens* strain BT-1 by 128 combinations of NCR peptides (1 mg/ml) in BM7 medium. Values along the top of a plot indicate the 96-well plate number the growth inhibition assay was performed in. The red line (“Negative_Control” in legend) in the plot is the growth of *L. crescens* strain BT-1 in the absence of inhibitors. The positive control (black lines with squares) represent *L. crescens* BT-1 growth in the presence of the antimicrobial polymyxin B sulfate at 0.1 mg/ml. The ‘Blank’ treatment contains BM7 medium only. Note the difference in x-axis values in panels Plate 1, 4, 5, and 6.

### Multiplexing peptides in the bacterial culture assays showed no positive, synergistic effects

After the initial screening of the 183 NCR peptides, we examined the possibility of growth inhibition synergy among the top ten most inhibitory NCR peptides (**Table S3**). To prevent having to test every possible pairwise combination, we performed a resolution five fractional factorial experimental design to efficiently evaluate many NCR peptide pairs simultaneously. A resolution five fractional factorial assay prevents confounding two-factor interactions with their main effects, enabling the detection of statistically significant interactions when they are indeed present. We utilized the ‘FrF2’ R package to create the fractional factorial experimental design. The FrF2 package uses graph theory algorithms to select pairs of treatments (i.e., NCR peptides) that are orthogonal (one peptide not overrepresented compared to another peptide) and independent of one another. Hence, instead of testing 1,024 pairwise combinations individually, the fractional factorial design streamlined the experiment to only 128 combinations of ten NCR peptides to be screened simultaneously (**Fig. 3**). Of the ten NCR peptides examined, peptides 803626 and 803627 were the only peptides with a significant effect on the growth inhibition (**Fig. S3, S4**). Of note, the interaction resulted in a decrease in growth inhibition (**Fig. S3**).

**Figure 3.**
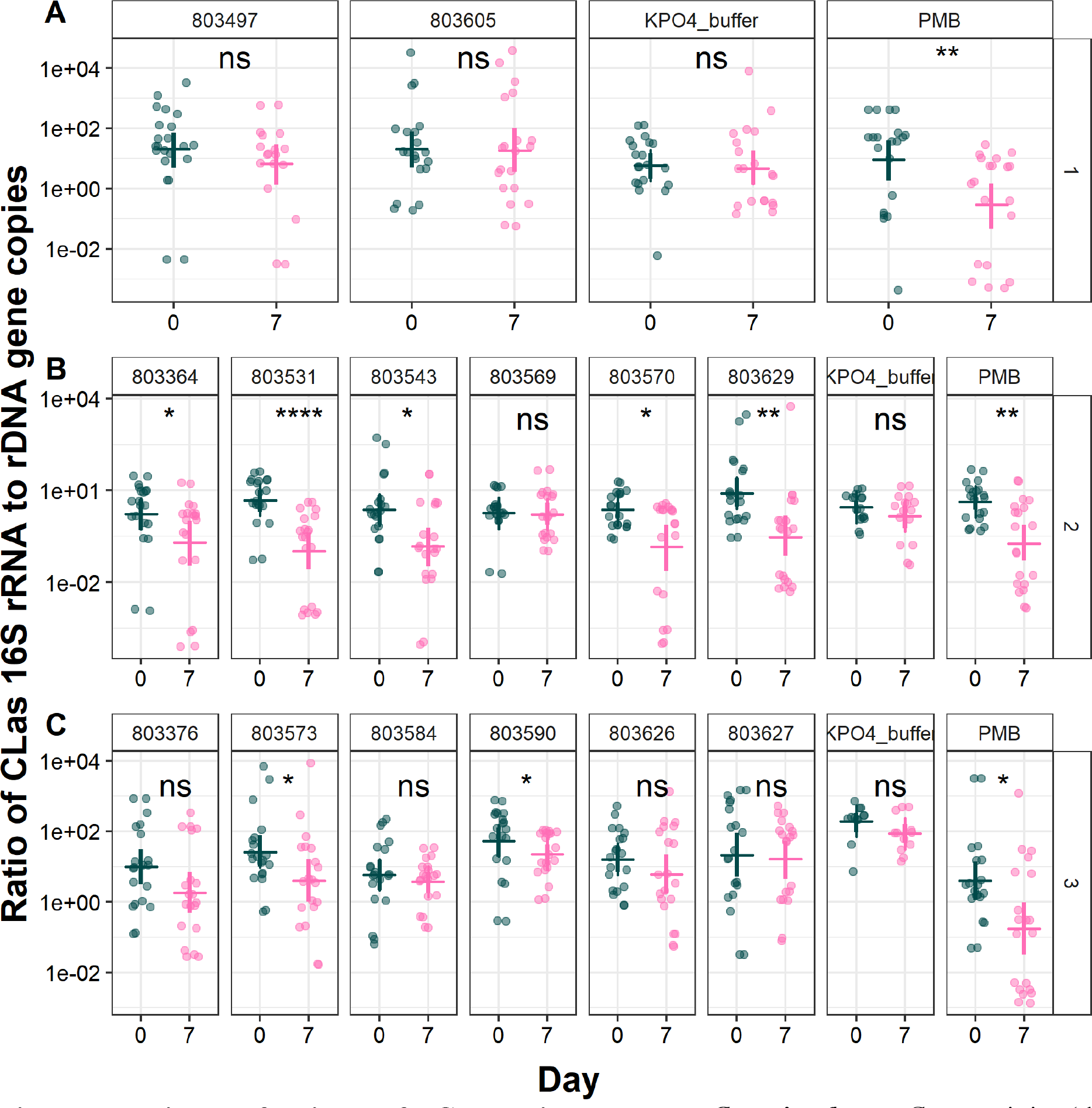
Excised leaf delivery of NCR peptides to target *C*Las *in planta*. *C*Las activity (the ratio of CLas 16S rRNA to rDNA gene copies) measured in detached *Citrus medica* (citron) leaves at 0 and 7 days following exposure to 0.1 mM potassium phosphate buffer (pH 5.8; label KPO4_buffer), NCR peptides (label 803XXX), or the antimicrobial peptide polymyxin B sulfate (label PMB in figure). Each point represents a technical duplicate qPCR result from the DNA and RNA extracted from leaves used in each treatment. The “ns” in figure panels indicates a non-significant Wilcoxon rank sum test result at α = 0.05. Asterisks indicate significance of Wilcoxon rank sum tests at p ≤ 0.05 (*), p ≤ 0.01 (**), p ≤ 0.001 (***), and p ≤ 0.0001 (****). Cross bars and vertical lines in each panel represent the nonparametric bootstrapped population mean and 95% confidence intervals (1,000 bootstraps; “mean_cl_boot” option in R stat_summary() function), respectively. A, B and C represent the three independent experiments that were performed to generate the final plot.

### Excised leaf delivery of NCR peptide 20-mers suggests NCR peptides have diverse modes of action against *C*Las

The 14 top-performing NCRs from the bacterial culture assays (**Table S3**) were selected for testing in an excised leaf assay to assess their effect on an established *C*Las infection. In these experiments, excised citrus leaves infected with *C*Las (validated by qPCR) were removed from infected citrus trees for treatment with the NCRs. The antimicrobial peptide Polymyxin B (PMB) was also tested. To our knowledge, PMB has not yet been reported in the literature to move in plant phloem or to inhibit *C*Las, but given its excellent performance as an antimicrobial peptide in the bacterial growth assays, we included PMB as a control in all experiments. After seven days of treatment, the leaves were analyzed using RT-qPCR or qPCR to measure the total number of copies of *C*Las rRNA and rDNA they contained, respectively. *C*Las activity was estimated from the ratio *C*Las rRNA to *C*Las rDNA gene copies (CLas rRNA:CLas rDNA), where a lower value for the ratio indicates higher antibacterial activity. The excised leaf assay revealed that 7/14 (50%) of the NCR peptides tested resulted in a significant reduction in the *C*Las 16S transcript-to-gene ratio over a seven-day incubation period, suggesting robust inhibition of CLas *in planta* by the NCR peptides (**Table 2**, **Fig. 3**).

**Table 2.**
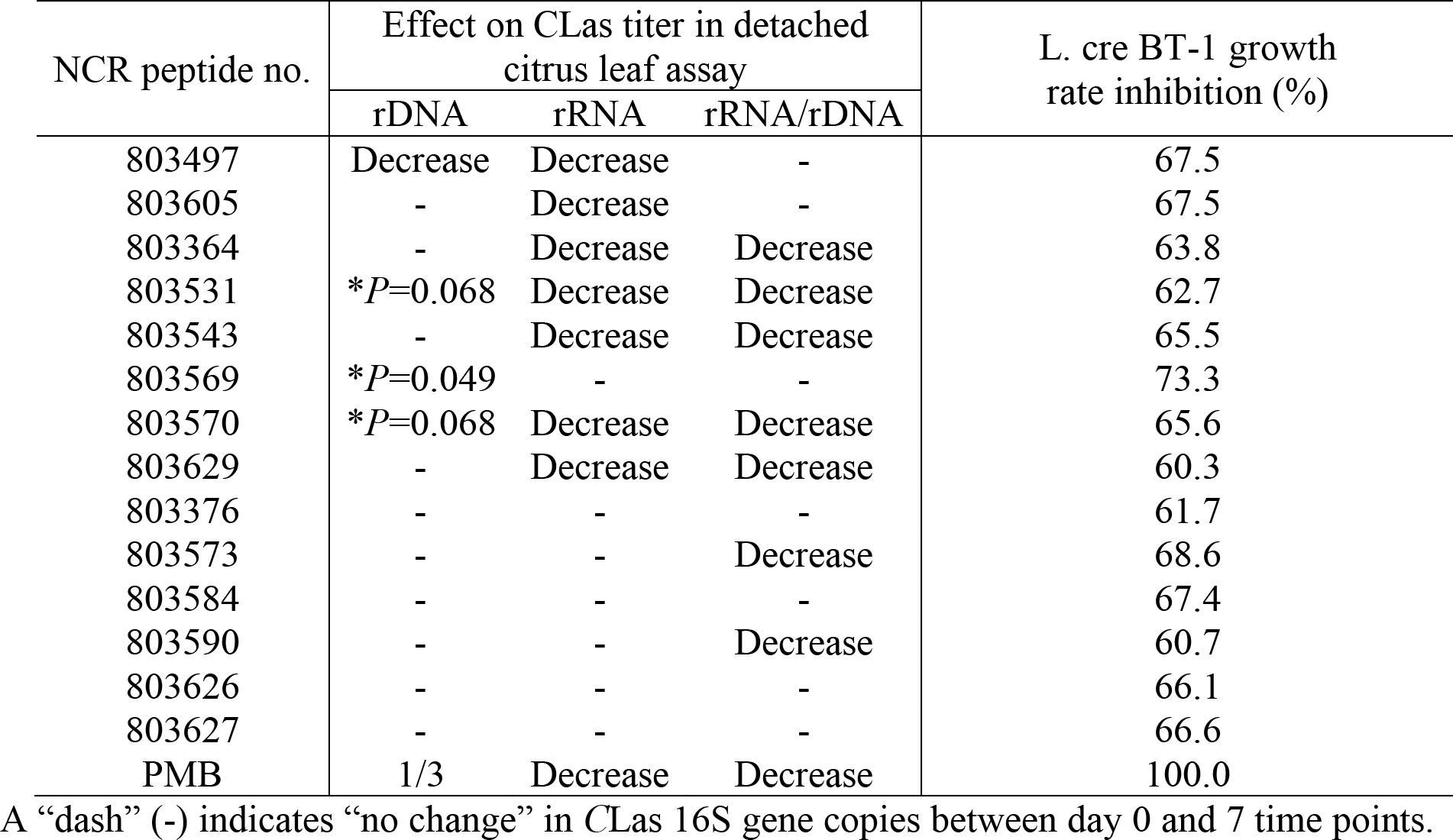
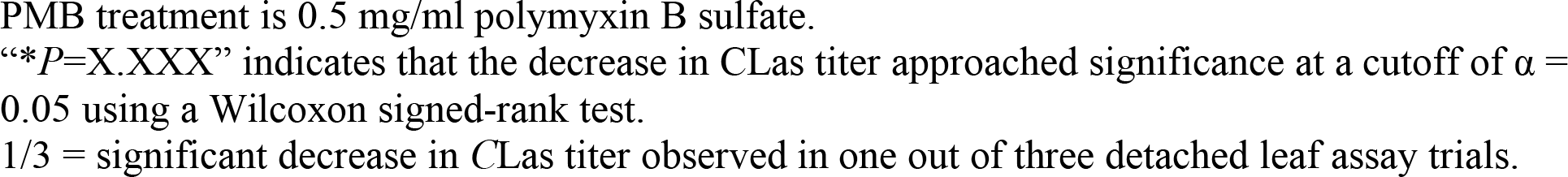
Effect of the top 14 NCR peptides (1 mg/ml) and PMB (0.5 mg/ml) on ‘*Candidatus* Liberibacter asiaticus’ 16S rRNA, rDNA, or rRNA/rDNA ratio in citrus detached leaf assays and their effect on *L. crescens* strain BT-1 growth rate.

| NCR peptide no. | Effect on CLas titer in detached citrus leaf assay |  |  | L. cre BT-1 growth rate inhibition (%) |
| --- | --- | --- | --- | --- |
|  | rDNA | rRNA | rRNA/rDNA |  |
| 803497 | Decrease | Decrease | - | 67.5 |
| 803605 | - | Decrease | - | 67.5 |
| 803364 | - | Decrease | Decrease | 63.8 |
| 803531 | * *P* =0.068 | Decrease | Decrease | 62.7 |
| 803543 | - | Decrease | Decrease | 65.5 |
| 803569 | * *P* =0.049 | - | - | 73.3 |
| 803570 | * *P* =0.068 | Decrease | Decrease | 65.6 |
| 803629 | - | Decrease | Decrease | 60.3 |
| 803376 | - | - | - | 61.7 |
| 803573 | - | - | Decrease | 68.6 |
| 803584 | - | - | - | 67.4 |
| 803590 | - | - | Decrease | 60.7 |
| 803626 | - | - | - | 66.1 |
| 803627 | - | - | - | 66.6 |
| PMB | 1/3 | Decrease | Decrease | 100.0 |

A “dash” (-) indicates “no change” in CLas 16S gene copies between day 0 and 7 time points.
PMB treatment is 0.5 mg/ml polymyxin B sulfate. “\*P=X.XXX” indicates that the decrease in CLas titer approached significance at a cutoff of $\alpha = 0.05$ using a Wilcoxon signed-rank test. 1/3 = significant decrease in CLas titer observed in one out of three detached leaf assay trials.

### Five NCR peptides prevent the development of *D. citri* adults with high titers of *C*Las

Acquisition of the *C*Las bacterium by psyllid nymphs is the first step required for tree-to-tree transmission. *C*Las cquisition is quantifiable in laboratory bioassays involving the development of psyllid nymphs on excised, infected citrus leaves (Igwe et al. 2021). Due to the labor intensiveness of the nymph acquisition assay, five of the top performing 14 NCR peptides with the lowest GRAVY scores (indicating higher solubility) were selected for this experiment: 803543, 803570, 803364, 803531, and 803629 (**Table S1**). After the duration of the 21-day assay, a total of 80, 84, 38, 15, 29, 147, and 148 adult insects were collected from the 803543, 803570, 803364, 803531, 803629, PMB and KPO4 treatments, respectively. While some degree of insect mortality occurred in all treatments during the assay, NCR peptides 803531 and 803629 resulted in significantly higher insect mortality as compared to the buffer control (**Fig. S5**). All five of the tested NCR peptides and the PMB control reduced *C*Las titer in the excised leaves during the course of the experiment (**Fig. 4A**, **Fig. S6**), although the differences were not statistically significant (**Table 3**, **Table S4**). In the control treatment, 42.68% of adult insects that developed on leaves supplied with buffer developed *C*Las titers of more than 10 cells per insect and 36.59% of the insects developed high titers, more than 100 *C*Las cells per insect (**Fig. 4B**, **Table 4**, **Table 5**, **Table S5**, **Fig. S6**, **Fig. S7**). The NCR peptides and PMB treatment prevented the development of high titer insects (**Fig. 4B**, **Table 4, Table S6**), with peptides 803543, 803364, 803531, and 803629 completely blocking the development of insects with more than 100 *C*Las cells (**Table 4**) and treatment of leaves with NCR peptide 803570 resulting in the development of only 2 insects with more than 100 *C*Las cells (**Table 4**).

**Figure 4.**
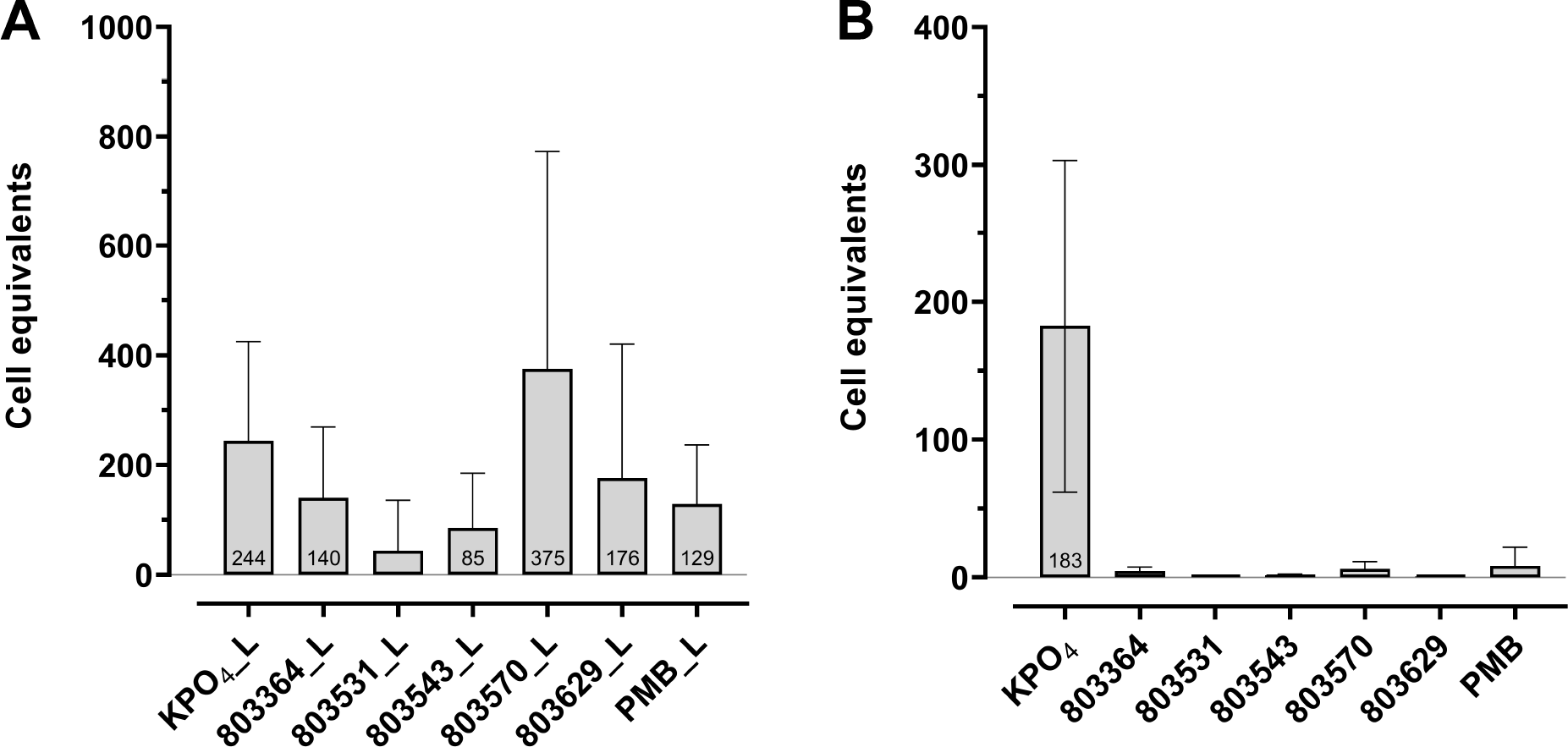
Treatment with five NCR peptides, 803364, 803531, 803543, 803570 and 803629 reduces *C*Las acquisition by *Diaphorina citri.* Plot of *C*Las cell equivalents per 50ng input DNA from excised citron leaves (A) and *Diaphorina citri* (B) samples involving treatments of the five top-performing NCR peptides against a known antimicrobial peptide polymyxin B sulfate (PMB) and buffer only control, potassium phosphate (KPO4). Titer values in (B) pertain to *C*Las titer in adult psyllids following treatment with the NCR peptides and controls using the excised leaf acquisition assay. In this iteration of the excised leaf assay (panel A), only 3531 showed significantly different *C*Las titer as compared to controls (*P =* 0.0288, See Table 3).

**Table 3.**
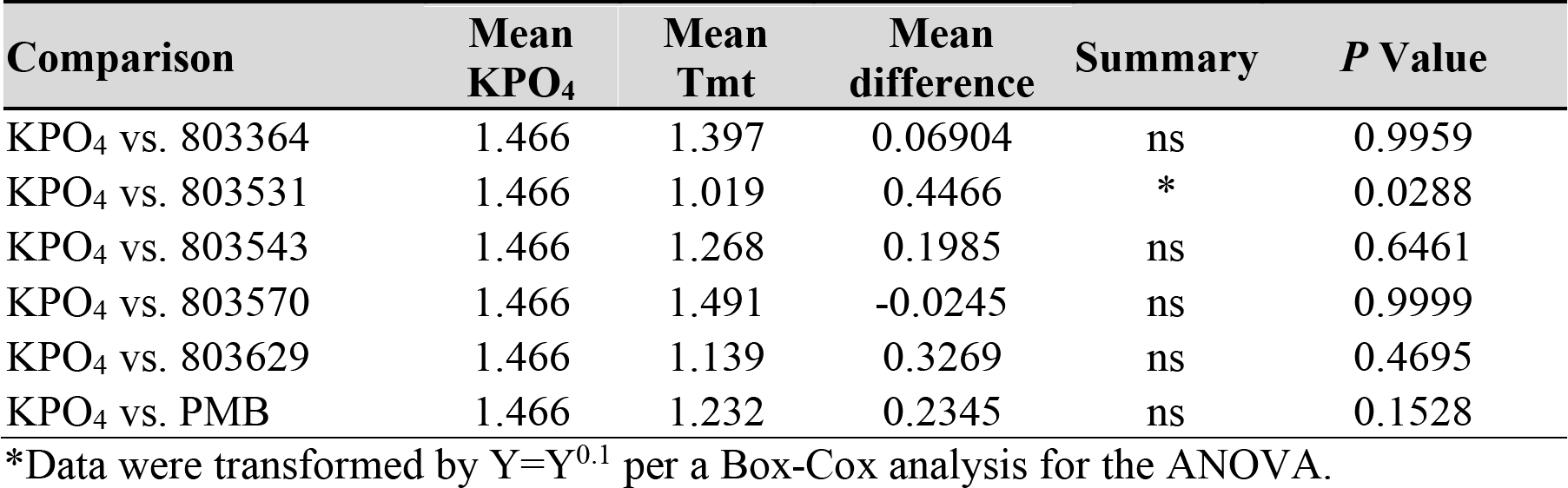
Welch’s ANOVA (*P* = 0.0537) on *C*Las cell equivalents* in leaves treated with NCR peptides used for the excised leaf acquisition assay followed by Dunnett’s T3 multiple comparison’s test (compares treatments against a control).

| Comparison | Mean KPO<sub>4</sub> | Mean Tmt | Mean difference | Summary | P Value |
| --- | --- | --- | --- | --- | --- |
| KPO<sub>4</sub> vs. 803364 | 1.466 | 1.397 | 0.06904 | ns | 0.9959 |
| KPO<sub>4</sub> vs. 803531 | 1.466 | 1.019 | 0.4466 | * | 0.0288 |
| KPO<sub>4</sub> vs. 803543 | 1.466 | 1.268 | 0.1985 | ns | 0.6461 |
| KPO<sub>4</sub> vs. 803570 | 1.466 | 1.491 | -0.0245 | ns | 0.9999 |
| KPO<sub>4</sub> vs. 803629 | 1.466 | 1.139 | 0.3269 | ns | 0.4695 |
| KPO<sub>4</sub> vs. PMB | 1.466 | 1.232 | 0.2345 | ns | 0.1528 |

\*Data were transformed by $Y=Y^{0.1}$ per a Box-Cox analysis for the ANOVA.

**Table 4:** Leaf delivery of five NCR peptides reduces the number of infective *Diaphorina citri* adults that developed on the treated leaves. No highly infected adults developed on the leaves treated with NCR peptides 803543, 803364, 803531 and 803629.

| Treatment | # leaves | Infected insects with<br>>100 CLas cells |  | Infected insects with<br>>10 CLas cells |  |
| --- | --- | --- | --- | --- | --- |
|  |  | No. | % | No. | % |
| NCR 803543 | 10 | 0/80 | 0 | 4/80 | 5 |
| NCR 803570 | 10 | 2/84 | 2.38 | 6/84 | 7.14 |
| NCR 803364 | 10 | 0/38 | 0 | 5/38 | 13.16 |
| NCR 803531 | 10 | 0/15 | 0 | 0/15 | 0 |
| NCR 803629 | 10 | 0/29 | 0 | 1/29 | 3.45 |
| PMB | 30 | 2/147 | 1.36 | 5/147 | 3.40 |
| KPO<sub>4</sub> | 30 | 34/148 | 22.97 | 55/148 | 37.16 |

Number of CLas cells >100 and >10 in the treatments and controls

**Table 5.**
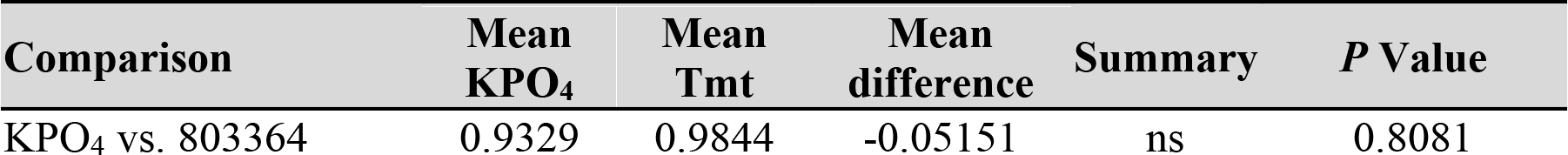
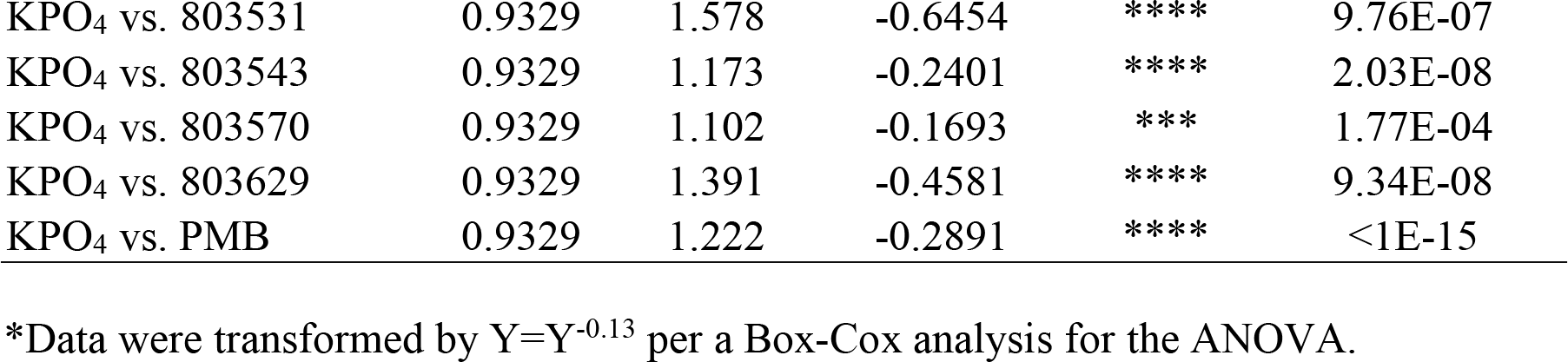
Welch’s ANOVA (p < 1E-15) on *C*Las cell equivalents* in *Diaphorina citri* individuals following acquisition from infected leaves followed by Dunnett’s T3 multiple comparison’s test.

| Comparison | Mean KPO<sub>4</sub> | Mean Tmt | Mean difference | Summary | P Value |
| --- | --- | --- | --- | --- | --- |
| KPO<sub>4</sub> vs. 803364 | 0.9329 | 0.9844 | -0.05151 | ns | 0.8081 |
| KPO<sub>4</sub> vs. 803531 | 0.9329 | 1.578 | -0.6454 | **** | 9.76E-07 |
| KPO<sub>4</sub> vs. 803543 | 0.9329 | 1.173 | -0.2401 | **** | 2.03E-08 |
| KPO<sub>4</sub> vs. 803570 | 0.9329 | 1.102 | -0.1693 | *** | 1.77E-04 |
| KPO<sub>4</sub> vs. 803629 | 0.9329 | 1.391 | -0.4581 | **** | 9.34E-08 |
| KPO<sub>4</sub> vs. PMB | 0.9329 | 1.222 | -0.2891 | **** | <1E-15 |

\*Data were transformed by $Y=Y^{-0.13}$ per a Box-Cox analysis for the ANOVA.

## Discussion

The HLB bacterium *C*Las and most fastidious Liberibacter spp. relatives are unculturable. It is not practical to screen a large number of antimicrobial compounds against *C*Las in leaves or insects (Kennedy et al. 2023). Therefore, we used the culturable relative of *C*Las, *L. crescens*, as a surrogate for *C*Las in bacterial culture assays, which can be carried out in 96-well plates, a more high-throughput way to narrow down the more than 600 NCR candidates to high priority candidates for testing against *C*Las in more laborious assays. NCR peptides were prioritized for synthesis and screening based on the peptide GRAVY score, with an emphasis on the most hydrophilic peptides. We predicted that small hydrophilic peptides would most likely be water soluble and phloem mobile. Furthermore, weak but positive correlations were observed between ability to inhibit *L. crescens* growth and peptide hydrophilicity and number of cystine residues. When screening pairwise combinations of NCRs in the bacterial culture assay, we observed additive activity for many pairs, but no synergistic increase in inhibition by the paired NCRs. In some cases, we observed suppression of growth inhibition with pairs of NCRs compared to individual NCR peptide treatments. Combinations that included either of two particular peptides consistently showed less activity than the individual peptides. While the reasons for this are not yet clear, the lesser effects of these peptides in combination may be due to their chemical properties, *e.g.*, lower solubility than the other 12 peptides or binding to the other peptide in the pairwise combination. The NCR peptides are diverse and are hypothesized to have a variety of biological roles in maintaining populations of the alphaproteobacterium *S. meliloti* in its leguminous host (Shabab et al. 2016), so the counteractive effect we have observed among some NCR peptides is not surprising. Moreover, their native expression is tightly spatially regulated, for example cell-type specific or in response to different conditions. Information regarding combinations of peptides that are counteractive is helpful in the development of field deployable formulations or expression in transgenic citrus. Since we only tested 10 NCR peptides for their ability to synergistically inhibit *L. crescens* strain BT-1 growth, it is possible other NCR peptides that were not top performers individually may perform better when combined with the most inhibitory peptides and could be examined in future research.

Analysis of NCR peptide treatment in plants in the excised leaf uptake experiments enabled us to determine both the antimicrobial activity and vascular mobility of the peptide. These are two required properties for antimicrobial peptides effective as HLB therapies. While the use of other powerful *C*Las screening systems have been reported, such as the hairy root system (Irigoyen et al. 2020; Mandadi et al. 2020), only assays that incorporate vascular mobility (movement in xylem or phloem) as a component of the screen, such as the excised leaf assay, can ascertain whether an antimicrobial peptide is mobile and stable in the citrus plant’s vascular system (Kennedy et al. 2023). In our experiments, we evaluated both *C*Las DNA and RNA since some antimicrobials may induce dormancy or initial death of the bacterium without rapid induction of cell lysis and DNA degradation. Thus, we posit that adverse effects on *C*Las may be more quickly observed by changes in the rRNA, which is consistent with the observations we report here. Evaluation of both nucleic acids allows a deeper understanding of how the *C*Las bacterium is responding to NCR treatment. The *in planta* screening produced variable results, and a large number of replicates are required to see an effect. The reasons for the variability may include the patchy distribution of *C*Las in each leaf (Fu et al. 2019), uneven distribution of the NCR peptides due to the vascular pathology induced by HLB (Ma et al. 2022), or the fact that only a single dose of NCR peptide was applied to each excised leaf during the screen. Nonetheless, the screen we performed here was sufficient for us to select peptides with some activity for further evaluation in our nymph acquisition assay.

Interestly, *C*Las titers were significantly lower during the excised leaf assay following NCR peptide treatment (Fig. 3) but not in the NCR-treated leaves following the nymph acquisition assay (Fig. 4), suggesting that psyllid feeding may elevate *C*Las titer in infected leaves. Analysis of our lab colonies shows that psyllid feeding increases *C*Las titer in the leaf (data not shown), which may explain the difference obsered, but there are other possibilities. The prolonged incubation required in the nymph acquisition assay may have reduced the effectiveness of the NCR peptide. Alternatively, it is possible that the NCR peptides interfered with *C*Las acquisition by psyllid nymphs due to induction of changes in *C*Las physiology or morphotype in the plant and not necessarily by killing *C*Las, so populations of *C*Las could remain viable and rebounded despite being unable to be acquired by psyllids. Differentiating between live and dead *C*Las extracted from plant tissue is not reliable (reviewed by (Kennedy et al. 2023), so this was not attempted. NCR peptides have been shown to regulate the morphotype of *S. meliloti* (Montiel et al. 2017), including induction of biofilm formation and production of extracellular vesicles. One hypothesis is that the NCR peptides modified *C*Las morphology in plants so as to block insect acquisition. Whole-tree experiments will be required to determine whether NCR peptide treatments would have similar effects when psyllids acquire *C*Las from flush tissue. Since *C*Las acquisition and transmission rates are highly correlated (Ammar et al. 2018) and almost zero *C*Las cells were acquired by psyllids during the NCR peptide treatments, transmission assays were not necessary to perform. The results from the nymph acquisition assay demonstrates the power of using insects as the indicator for antimicrobial peptide treatment efficacy, as the insects will only acquire live bacteria in a morphotype suitable or capable of replicating in insect tissues.

HLB represents a devastating, recalcitrant pathosystem that has resulted in significant economic losses for the United States citrus industry. The NCR peptides identified in our screen may be developed for deployment in the field either through direct application to trees, incorporation into a cross-protective synthetic bacterial or viral strains, or transgenic plant expression to control *C*Las. Our work adds to the body of literature that shows the promise of peptide therapies for treatment of HLB (Huang et al. 2021; Wang 2021). Our results show that NCR peptides offer an unexplored, protective strategy that could salvage the rapidly declining U.S. citrus industry if peptide delivery strategies were mature enough to deploy the peptides to trees in the groves. Since these are plant-derived peptides, they may be registered by the United States Environmental Protection Agency as biopesticides and face fewer regulatory hurdles during commercialization. It is plausible that the NCR peptides we identified as having killing- activity against *C*Las may also be effective at inhibiting the growth and/or transmission of other bacterial species threatening the citrus industry, such as *Spiroplasma citri*, the phloem-limited bacterium that causes citrus stubborn disease. Peptides targeting the relationship among *C*Las, the plant and the insect vector will be the foundation for developing a field-deployable control strategy for HLB, which is vitally needed for the maintenance of citrus production in the US and worldwide. In particular, the five NCR peptides reported here to block the development of highly infective psyllids could be crucial for deployment in California to protect healthy citrus trees from becoming infected because the disease is not yet endemic in commercial production areas and in endemic areas, such as Florida, to prevent *C*Las spread to new plantings.

## Acknowledgements

The authors are grateful to Luke Thompson (USDA ARS), Julie Blaha (Cornell), Amani Helo (Cornell), and Chad Thomas (Cornell) for their assistance with maintaining citrus plants and insect colonies. We thank the California Citrus Research Board for funding to support this research through Project # 5300-202. The work was also supported by a cooperative research and development agreement between AgroSource, Inc. and USDA ARS and USDA ARS Project #s: 8062-22410-007-000-D and 6034-21000-018-000-D. We are grateful to Dr. Robert Krueger, Curator of the National Clonal germplasm Repository for Citrus at the USDA ARS in Riverside, CA, for providing clean seeds to our lab. We thank Dr. Mark Trimmer (AgroSource, Inc) and members of the Heck Lab for helpful discussion and feedback on the manuscript.

## Supplementary Information

**Figure S1.**
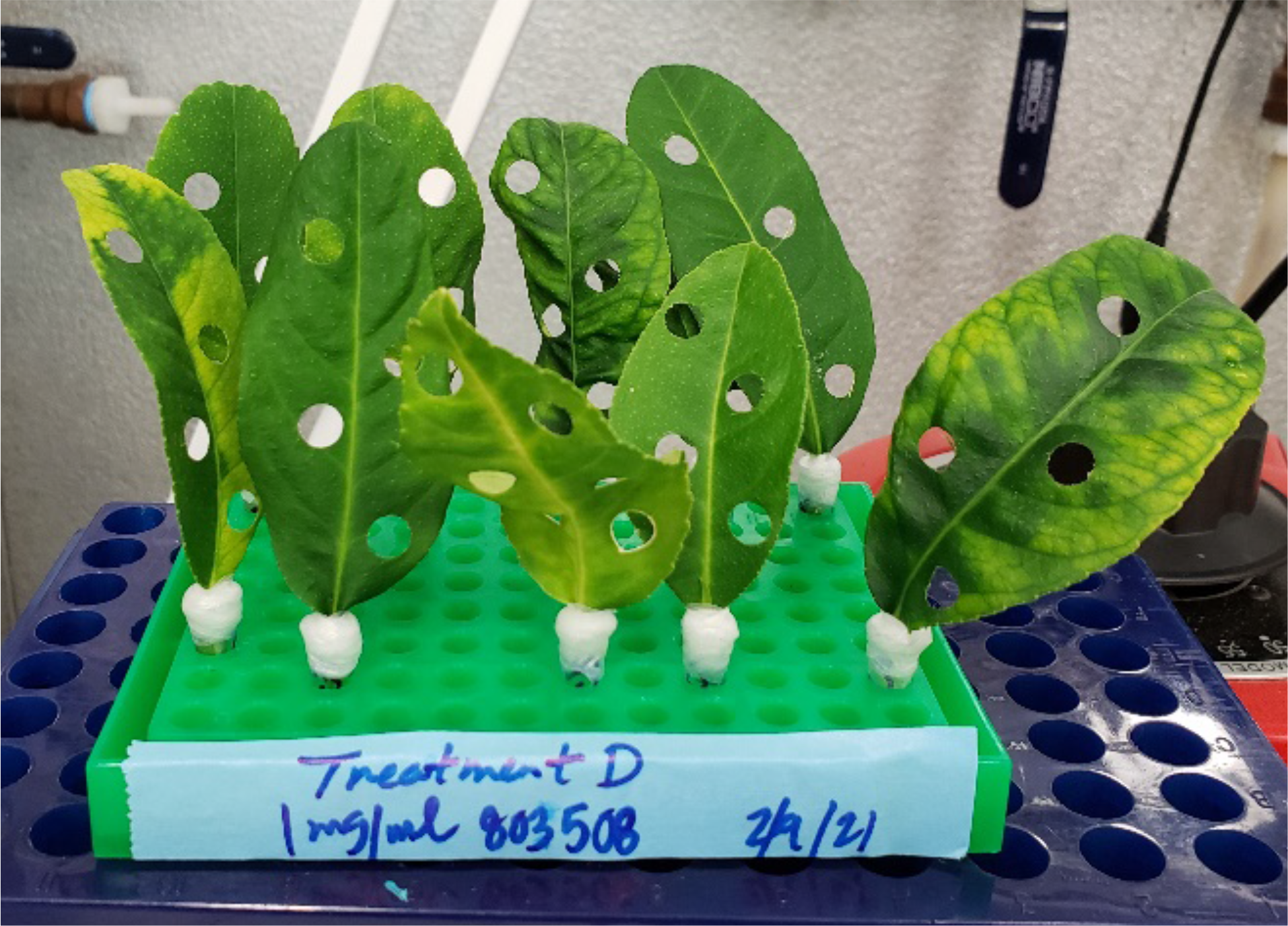
Example of citrus excised leaf assay. The ten citrus leaves above were confirmed to be infected by ‘*Candidatus* Liberibacter asiaticus’ (*C*Las) and are being treated with the NCR peptide 803508 in this experiment.

**Figure S2.**
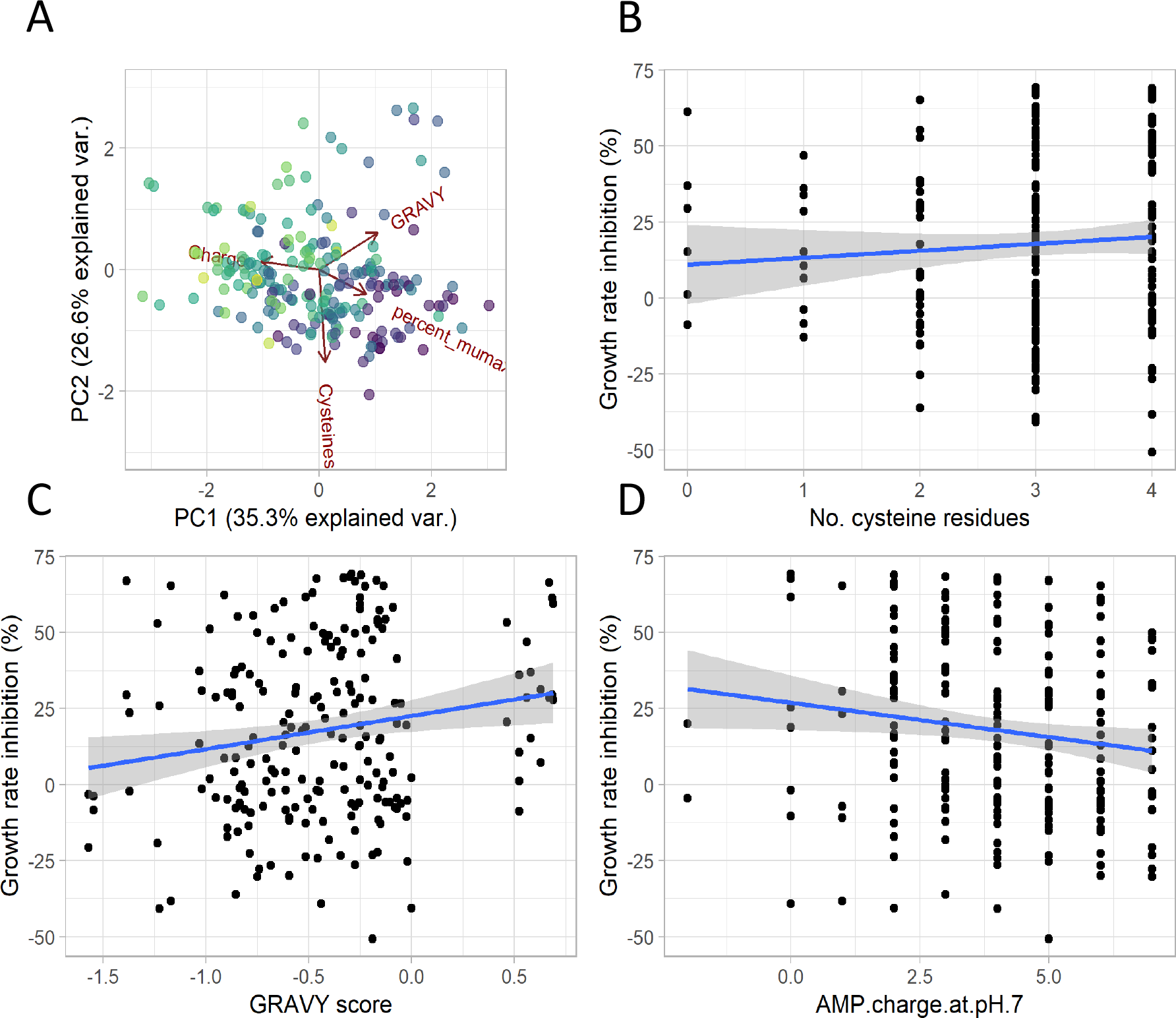
Growth rate inhibition as a function of the physiochemical properties of the NCR peptides. A. Principal components analysis of the relationship between percent growth rate inhibition (percent_mumax) and select physiochemical properties of each NCR peptide. The GRAVY score, antimicrobial peptide charge at pH 7, and number of cysteine residues displayed some level of correlation with percentage of growth rate inhibition by the NCR peptides examined (darker colors indicate a greater percentage of growth rate inhibition with a range of 0 to 73.3%). B. Regression on number of cysteine resides as a function of growth rate inhibition of *L. crescens.* C. GRAVY score as a function of growth rate inhibition. D. Predicted NCR peptide charge at pH 7 as a function of growth rate inhibition. Spearman rank correlations for plots B, C, and D were 0.15, 0.09, and -0.15 and were only significant for B, and D (p values 0.025 and 0.026, respectively).

**Figure S3.**
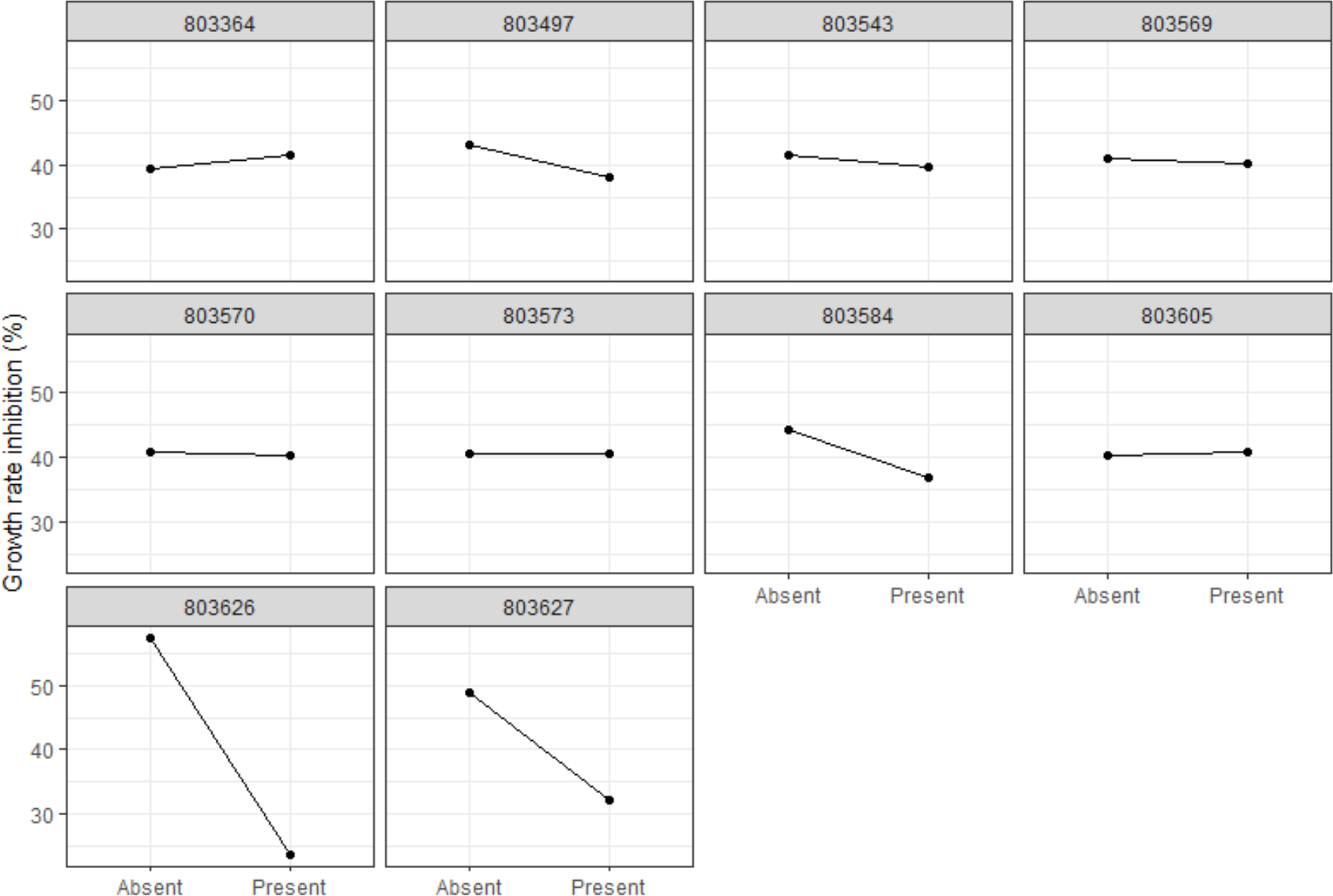
Main effects plot of the NCR peptide fractional factorial design. No statistically significant increases in growth inhibition were identified, and two peptides (803626 and 803627) were identified whose presence significantly reduced growth inhibition.

**Figure S4.**
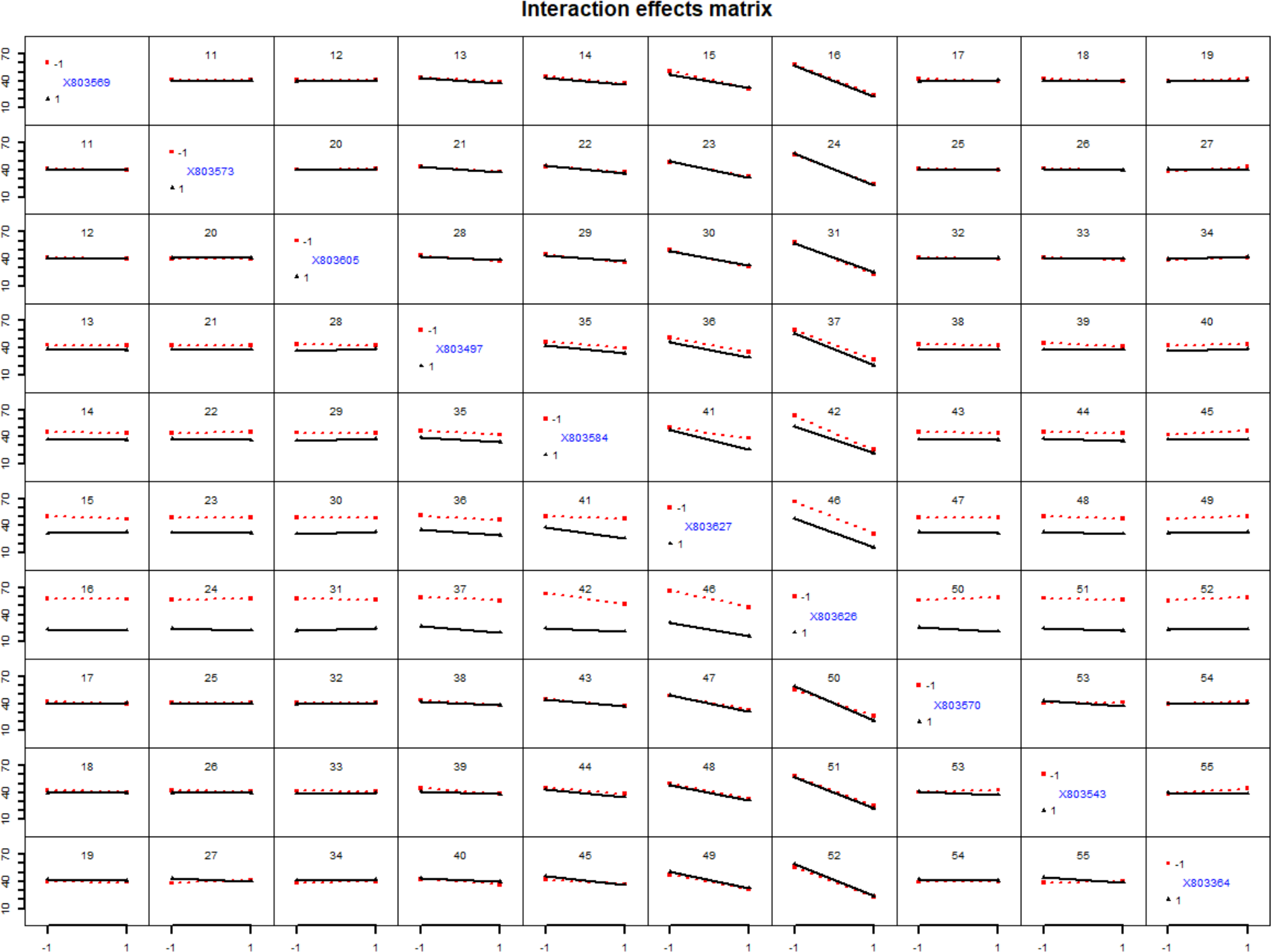
Interaction effects plot of the NCR peptide fractional factorial design. The y-axis is growth inhibition (%) compared to the control (*L. crescens* BT-1 in BM7 medium without inhibitor), and the x-axis is the presence or absence of each NCR peptide (blue labels inside plot are NCR peptide IDs). The only significant interactions identified among peptides was a decrease in growth inhibition (y -axis) when NCR peptides 803627 and 803626 were present. Otherwise, only non-significant interactions were identified.

**Figure S5:**
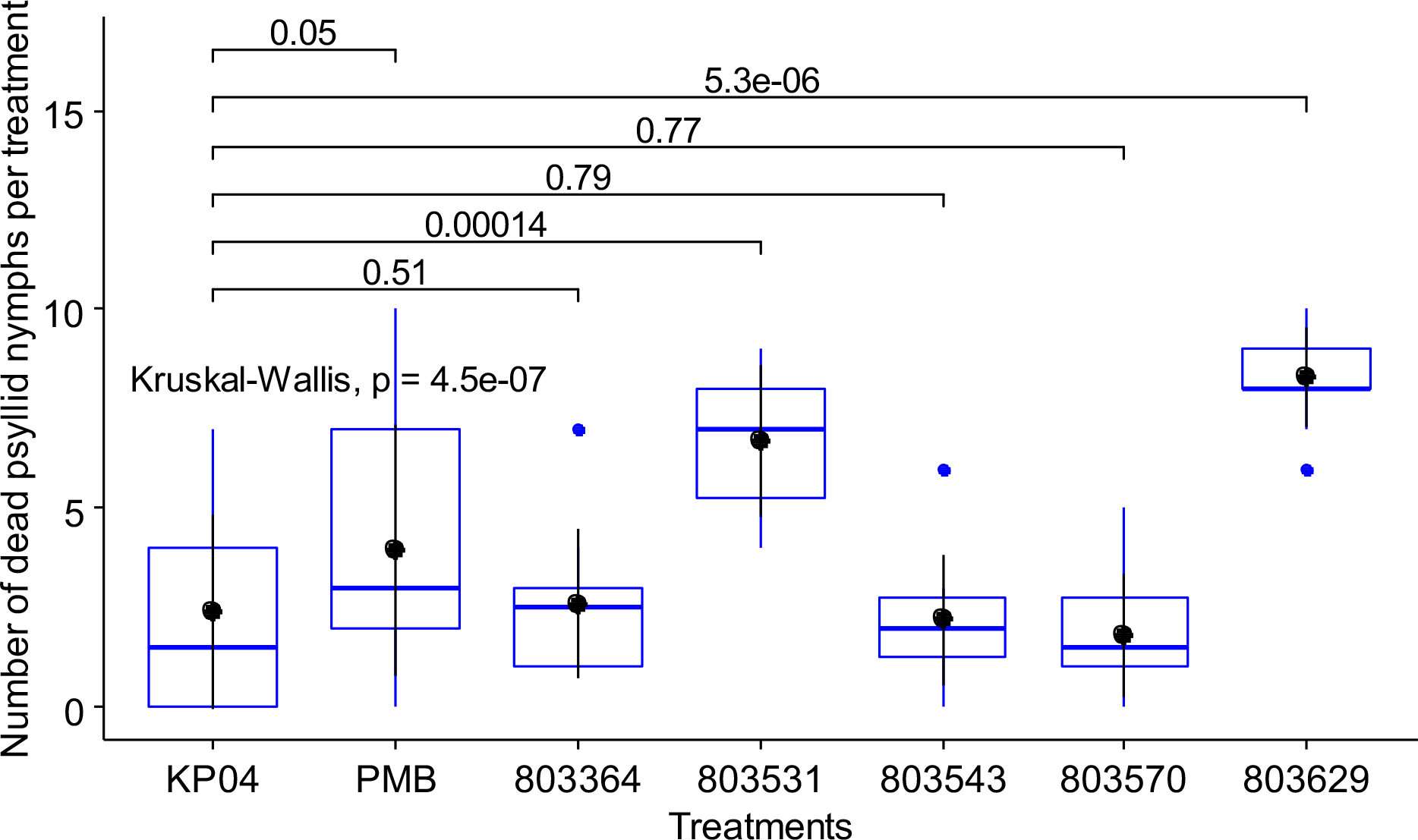
Treatment of excised citrus leaves with NCR peptides 803531 and 803629 induce mortality of nymph and adult *Diaphorina citri* in the nymph acquisition assay as compared to the potassium phosphate (KPO4) buffer control but NCR peptides 803364, 803543, 803570 or polymyxin B sulfate (PMB).

**Figure S6.**
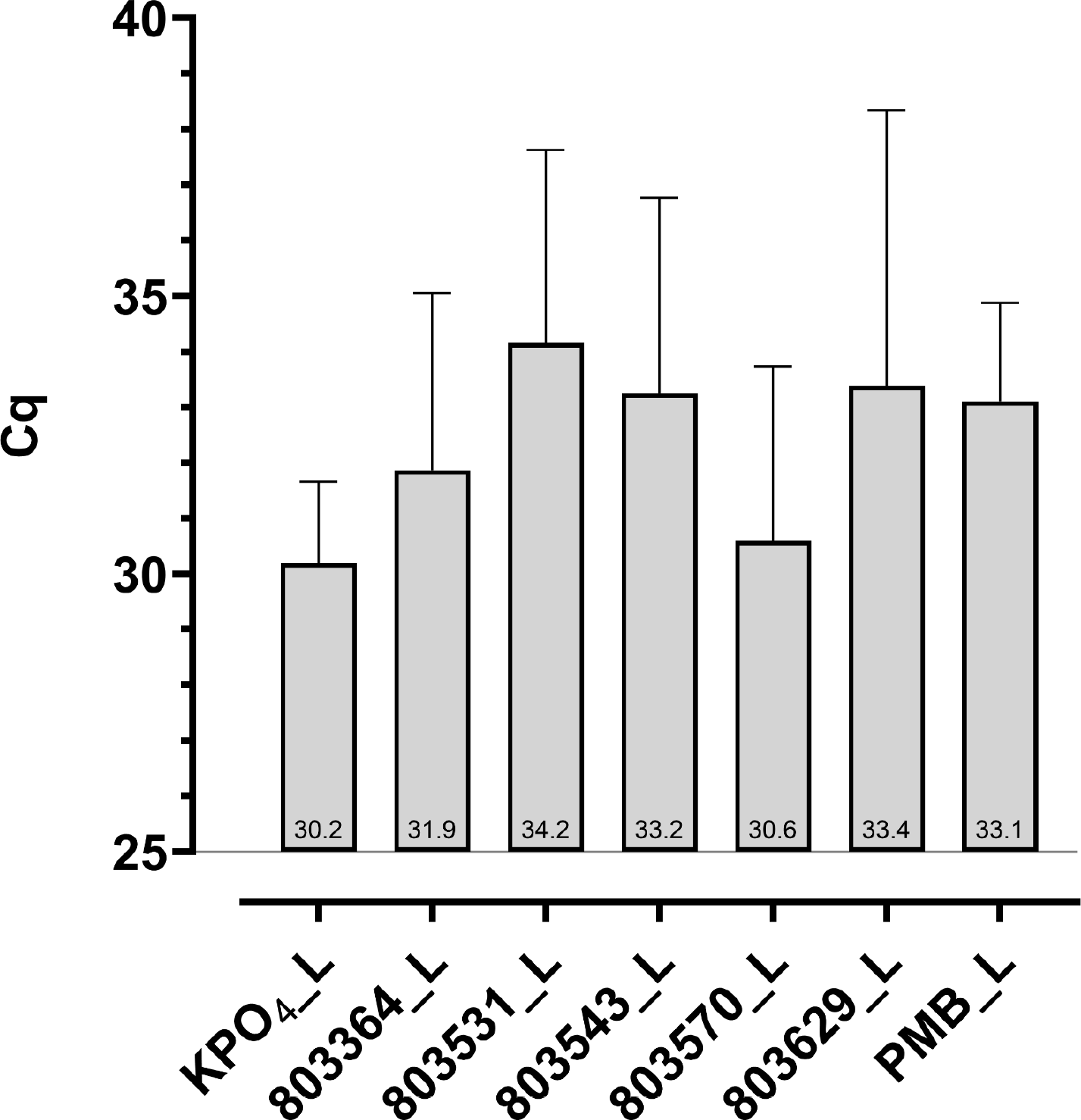
Impacts on *C*Las Cq values in *Citrus medica* leaf tissues following treatment with five NCR peptides, 803364, 803531, 803543, 803570 and 803629, a known antimicrobial peptide polymyxin B sulfate (PMB) and buffer only control, potassium phosphate (KPO4) using excised leaf delivery method. These are the Cq values of the leaf tissues that were used for the excised leaf acquisition assay. Mean Cq values in the leaf disks indicated at the bottom of each treatment bar.

**Figure S7.**
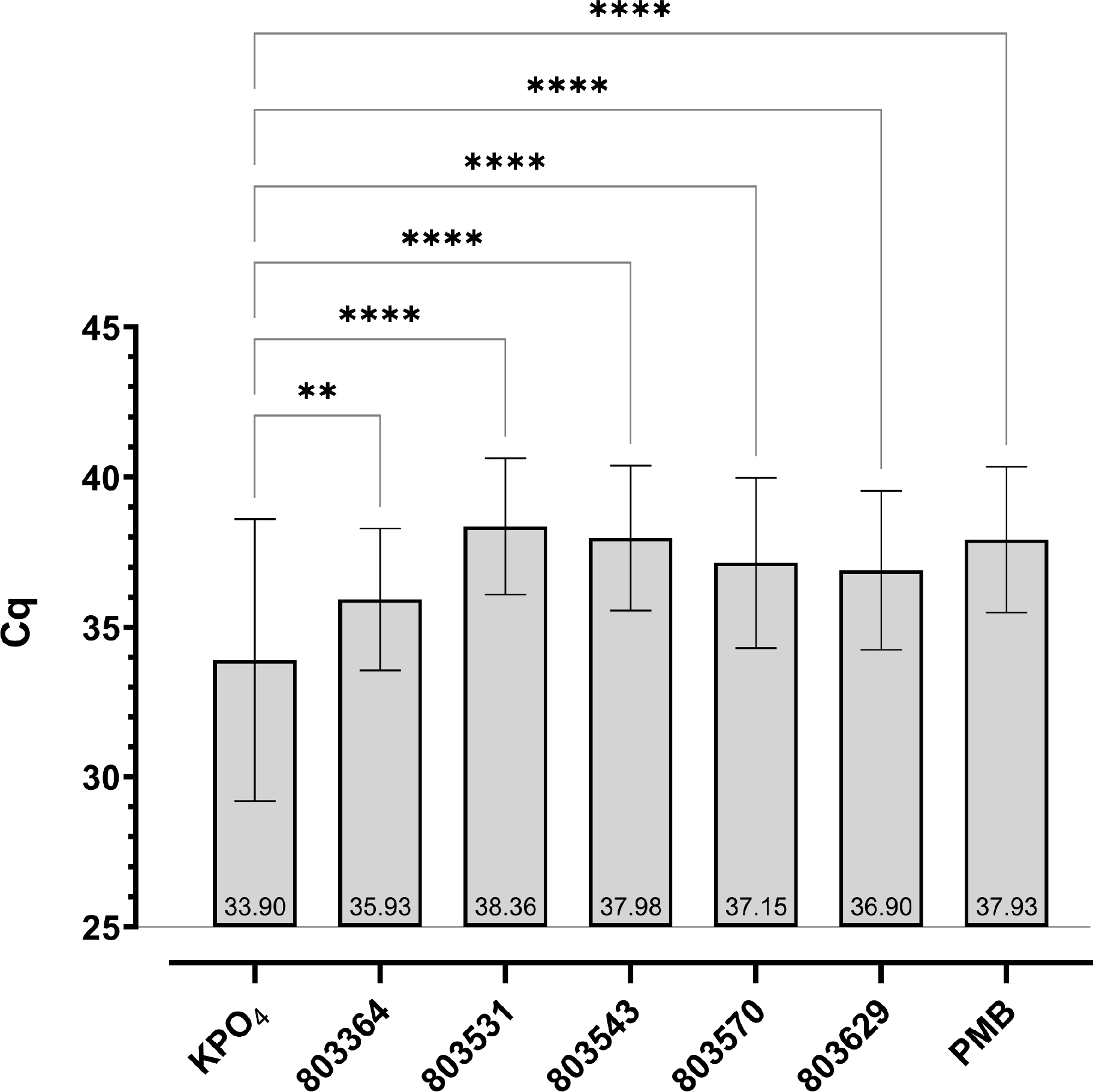
Impacts on *C*Las Cq values in adult *Diaphorina citri* following treatment with five NCR peptides, 803364, 803531, 803543, 803570 and 803629, a known antimicrobial peptide polymyxin B sulfate (PMB) and buffer only control, potassium phosphate (KPO4) using excised leaf delivery method. These are the Cq values of the insects that were used for the excised leaf acquisition assay. Mean Cq values in the leaf disks indicated at the bottom of each treatment bar.

Table S1. All predicted NCR peptides and GRAVY scores.

Table S2. The qPCR standard curve information used to calculate gene copy number reported in the manuscript.

Table S3. Top 14 NCR peptides, GRAVY scores, # cysteine residues, AMP charge at ph7 and %mumax.

**Table S4.**
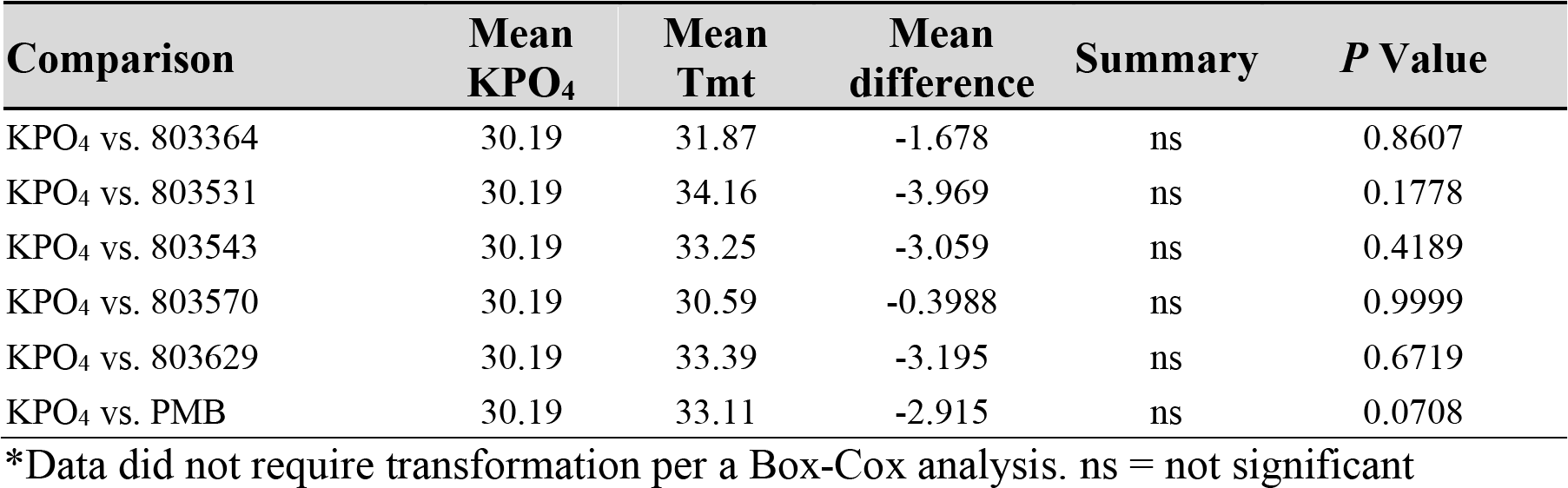
Welch’s ANOVA (p = 0.2970) on Cq of leaf disks from leaves used in the excised leaf acquisition assay followed by Dunnett’s T3 multiple comparison’s test.

**Table S5.**
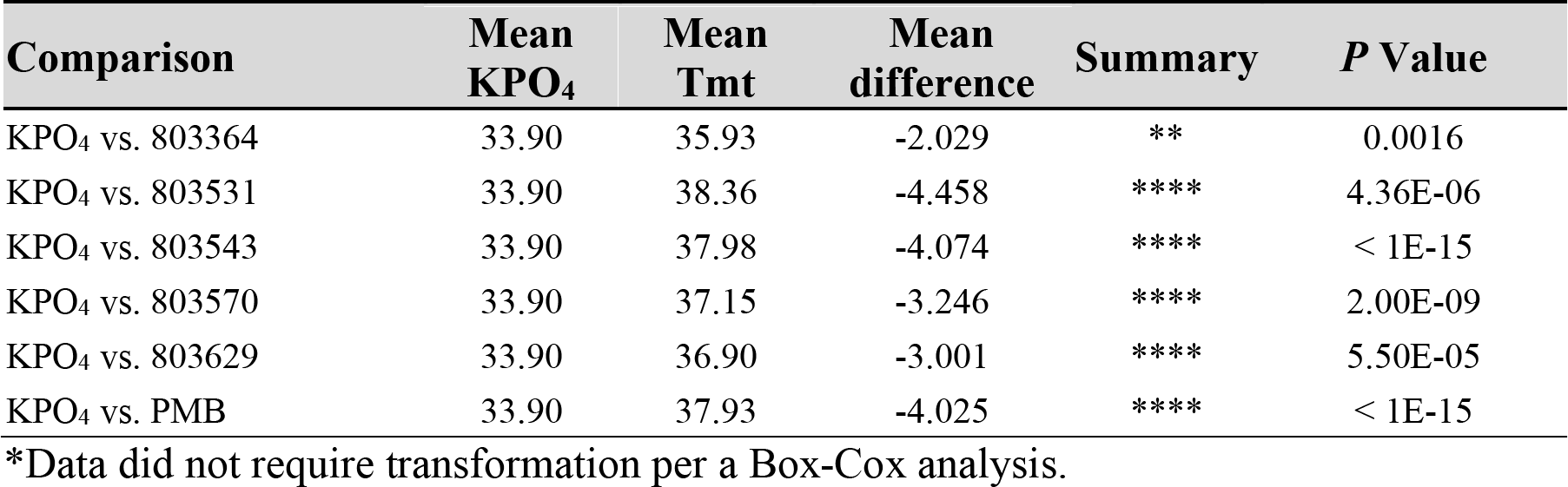
Welch’s ANOVA (p < 1E-15) on Cq of psyllid samples following NCR peptide treatment via excised leaves followed by Dunnett’s T3 multiple comparison’s test.

**Table S6.**
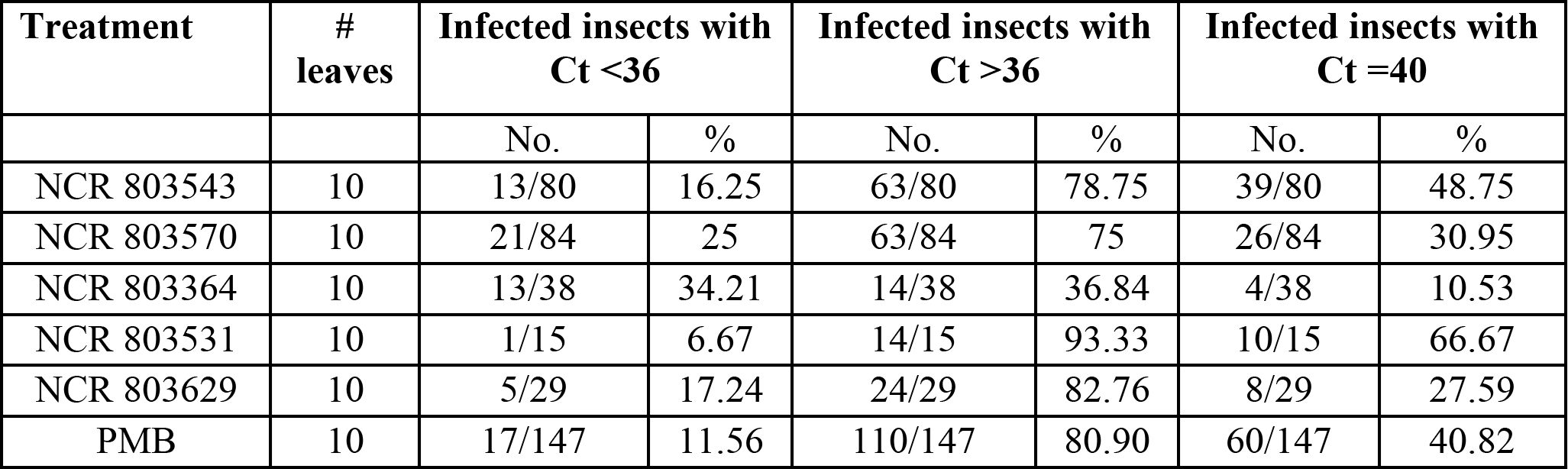
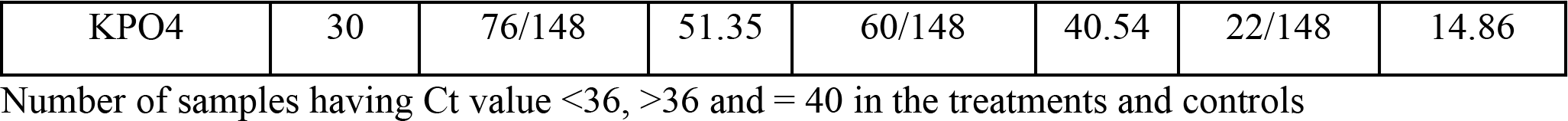
Delivery of five NCR peptides to *C*Las-infected citrus leaves reduces the number of infective *Diaphorina citri* that develop.

