## Supplementary material for "Plant-derived, nodule-specific cysteine rich peptides inhibit growth and psyllid acquisition of ‘*Candidatus* Liberibacter asiaticus’, the citrus Huanglongbing bacterium": Table S4

**Table S4.** Welch’s ANOVA (p = 0.2970) on Cq of leaf disks from leaves used in the excised leaf acquisition assay followed by Dunnett’s T3 multiple comparison’s test.

| **Comparison** | **Mean KPO_4_** | **Mean Tmt** | **Mean difference** | **Summary** | ***P* Value** |
| --- | --- | --- | --- | --- | --- |
| KPO_4_ vs. 803364 | 30.19 | 31.87 | -1.678 | ns | 0.8607 |
| KPO_4_ vs. 803531 | 30.19 | 34.16 | -3.969 | ns | 0.1778 |
| KPO_4_ vs. 803543 | 30.19 | 33.25 | -3.059 | ns | 0.4189 |
| KPO_4_ vs. 803570 | 30.19 | 30.59 | -0.3988 | ns | 0.9999 |
| KPO_4_ vs. 803629 | 30.19 | 33.39 | -3.195 | ns | 0.6719 |
| KPO_4_ vs. PMB | 30.19 | 33.11 | -2.915 | ns | 0.0708 |

*Data did not require transformation per a Box-Cox analysis. ns = not significant
