## Supplementary material for "Plant-derived, nodule-specific cysteine rich peptides inhibit growth and psyllid acquisition of ‘*Candidatus* Liberibacter asiaticus’, the citrus Huanglongbing bacterium": Table S5

**Table S5.** Welch’s ANOVA (p < 1E-15) on Cq of psyllid samples following NCR peptide treatment via excised leaves followed by Dunnett’s T3 multiple comparison’s test.

| **Comparison** | **Mean KPO_4_** | **Mean Tmt** | **Mean difference** | **Summary** | ***P* Value** |
| --- | --- | --- | --- | --- | --- |
| KPO_4_ vs. 803364 | 33.90 | 35.93 | -2.029 | ** | 0.0016 |
| KPO_4_ vs. 803531 | 33.90 | 38.36 | -4.458 | **** | 4.36E-06 |
| KPO_4_ vs. 803543 | 33.90 | 37.98 | -4.074 | **** | < 1E-15 |
| KPO_4_ vs. 803570 | 33.90 | 37.15 | -3.246 | **** | 2.00E-09 |
| KPO_4_ vs. 803629 | 33.90 | 36.90 | -3.001 | **** | 5.50E-05 |
| KPO_4_ vs. PMB | 33.90 | 37.93 | -4.025 | **** | < 1E-15 |

*Data did not require transformation per a Box-Cox analysis.
