## Supplementary material for "Plant-derived, nodule-specific cysteine rich peptides inhibit growth and psyllid acquisition of ‘*Candidatus* Liberibacter asiaticus’, the citrus Huanglongbing bacterium": Table S6

**Table S6.** Delivery of five NCR peptides to CLas-infected citrus leaves reduces the number of infective Diaphorina citri that develop.

| **Treatment** | **# leaves** | **Infected insects with**  **Ct <36** | | **Infected insects with**  **Ct >36** | | **Infected insects with**  **Ct =40** | |
| --- | --- | --- | --- | --- | --- | --- | --- |
|  |  | No. | % | No. | % | No. | % |
| NCR 803543 | 10 | 13/80 | 16.25 | 63/80 | 78.75 | 39/80 | 48.75 |
| NCR 803570 | 10 | 21/84 | 25 | 63/84 | 75 | 26/84 | 30.95 |
| NCR 803364 | 10 | 13/38 | 34.21 | 14/38 | 36.84 | 4/38 | 10.53 |
| NCR 803531 | 10 | 1/15 | 6.67 | 14/15 | 93.33 | 10/15 | 66.67 |
| NCR 803629 | 10 | 5/29 | 17.24 | 24/29 | 82.76 | 8/29 | 27.59 |
| PMB | 10 | 17/147 | 11.56 | 110/147 | 80.90 | 60/147 | 40.82 |
| KPO4 | 30 | 76/148 | 51.35 | 60/148 | 40.54 | 22/148 | 14.86 |

Number of samples having Ct value <36, >36 and = 40 in the treatments and controls
